# The cGAS-STING pathway regulates microglial chemotaxis in genome instability

**DOI:** 10.1101/2023.08.25.554654

**Authors:** Emily J. Talbot, Lisha Joshi, Peter Thornton, Mahya Dezfouli, Kalliopi Tsafou, Michael Perkinton, Svetlana V. Khoronenkova

## Abstract

Defective DNA damage signalling and repair is a hallmark of age-related and genetic neurodegenerative disease. One mechanism implicated in disease progression is DNA damage-driven neuroinflammation, which is largely mediated by tissue-resident immune cells, microglia. Here, we utilise human microglia-like cell models of persistent DNA damage and ATM kinase deficiency to investigate how genome instability shapes microglial function. We demonstrate that upon DNA damage the cytosolic DNA sensing cGAS-STING axis drives chronic inflammation and a robust chemokine response, exemplified by production of CCL5 and CXCL10. Transcriptomic analyses revealed that cell migratory pathways were highly enriched upon IFN-β treatment of human iPSC-derived microglia, indicating that the chemokine response to DNA damage mirrors type I interferon signalling. Furthermore, we find that STING deletion leads to a defect in microglial chemotaxis under basal conditions and upon ATM kinase loss. Overall, this work provides mechanistic insights into cGAS-STING-dependent neuroinflammatory mechanisms and consequences of genome instability in the central nervous system.

## INTRODUCTION

Accumulation of DNA damage in the central nervous system (CNS) is commonly observed in ageing and neurodegenerative diseases, including Alzheimer’s and amyotrophic lateral sclerosis^1–3^. Additionally, loss-of-function mutations in DNA damage response (DDR) genes underpin neurodegeneration in monogenic disorders, such as ataxia with oculomotor apraxia types 1-5 and ataxia-telangiectasia. Such vulnerability of the CNS to DDR defects is thought to stem from the interplay between cell intrinsic effects of DNA damage on post-mitotic neurons and glia as well as extrinsic signals resulting from glia-driven neuroinflammation^4,5^.

One mechanism linking genome instability and neuroinflammation is the cytosolic DNA sensing cGAS-STING pathway. Although responsible for mobilising antimicrobial responses to cytosolic pathogenic DNA during infection, the cGAS-STING pathway can mount innate immune responses to host DNA aberrantly localised in the cytosol following DNA damage^6^. The nucleotidyl transferase cGAS binds pathogenic double-stranded DNA (dsDNA) in the cytosol, catalysing synthesis of 2’,3’-cyclic GMP-AMP (cGAMP). cGAMP activates the immune adaptor stimulator of interferon genes (STING) at the endoplasmic reticulum (ER), inducing STING oligomerisation and translocation to the Golgi, where it recruits the TBK1 kinase and the interferon regulatory transcription factor IRF3. STING then acts as a platform for TBK1 autophosphorylation at S172, as well as phosphorylation of STING at S366 and IRF3 at S396, culminating in nuclear translocation of IRF3. STING signalling also activates NF-κB, leading to IRF3- and NF-κB-dependent transcription of type I interferons, interferon-stimulated genes (ISGs), and pro-inflammatory cytokines which mount the antimicrobial response^7,8^.

In the absence of infection, cytosolic accumulation of host DNA arising from genome instability can also activate STING-dependent autoimmunity. One such example is cGAS activation by nucleosome tethering at ruptured micronuclei, which can form during mitosis in cells harbouring unresolved DNA damage^9–11^. Moreover, cGAS is activated by nuclear dsDNA, RNA:DNA hybrids, telomeric DNA and mitochondrial DNA which can accumulate in the cytosol following genotoxic or mitochondrial stress^12–15^. Thus, cGAS-STING promotes inflammation in response to diverse self-DNA species.

In the brain, the strongest response to cGAS-STING activation occurs in microglia, the resident macrophages of the CNS^16^. Microglia maintain CNS homeostasis via surveillance of the microenvironment, phagocytosis of apoptotic cells and/or debris, synaptic pruning, and production of neurotrophic factors^17–20^. During infection or injury, microglia can transition towards a heterogenous ‘reactive’ state typified by enhanced phagocytosis, motility, proliferation, and production of inflammatory mediators and reactive oxygen/nitrogen species^21^. However, chronic microglial reactivity is implicated in neurodegenerative disease, including those driven by genome instability, such as ataxia-telangiectasia (A-T)^5,22^.

A-T is caused by loss-of-function mutations in ATM, a protein kinase which coordinates cellular responses to cytotoxic DNA double strand breaks (DSBs) and oxidative stress. More broadly, ATM preserves homeostasis by regulating redox balance, glucose metabolism, and immunity^23^. In humans, A-T manifests as cerebellar degeneration and ataxia, and microglia-driven neuroinflammation is increasingly implicated in disease phenotypes. We previously demonstrated that ATM loss in human microglia-like cells activates RELB/p52 NF-κB signalling and drives aberrant phagocytic activity, resulting in damage to the neuronal network *in vitro*^24^. Furthermore, microglial activation is observed in Atm-deficient murine microglia and correlates with motor neuron loss in the spinal cord of Atm-deficient rats^25–28^. Interestingly, cerebellar microglia exhibit characteristics which may predispose them to exacerbated immune responses to genome instability, including heightened immune vigilance and enhanced proliferation^29,30^. Indeed, gene signatures enriched in cytosolic DNA sensing, antiviral immunity, and proliferation are upregulated specifically in cerebellar microglia in A-T^31^. In addition, ATM deficiency activates cytosolic DNA sensing in diverse cell models, including primary murine microglia and A-T patient-derived brain organoids^6,26,32–35^. STING activation in ATM deficiency is also linked with neuronal damage via microglial secretion of IL-1β and inflammatory responses mediated by senescent astrocytes^26,32^. However, the molecular mechanisms of how STING signalling and type I interferon production shapes microglial function in genome instability remain unclear.

Here, we investigate how loss of ATM kinase and persistent DNA damage regulate microglial function. We find that genome instability drives chronic chemokine production in human microglia-like cells and the chemokine response is regulated by the cGAS-STING pathway. In agreement with this, bulk RNA-sequencing of iPSC-derived microglia treated with IFN-β implicated type I interferon signalling in the governance of microglial migration. Indeed, we find that STING deletion leads to defective microglial migration in basal conditions and upon loss of ATM function. We propose that STING-mediated chronic chemokine production may amplify neuroinflammation in genome instability via several different mechanisms, including the recruitment of peripheral immune cells to the CNS and excessive energy expenditure. Overall, this work provides mechanistic insight into STING-dependent regulation of microglial function upon persistent DNA damage.

## RESULTS

### Loss of ATM function in human microglia-derived cells drives STING pathway activation and a type I interferon response

Individuals with classical A-T phenotypes typically possess nonsense mutations leading to complete loss of functional protein, STING signalling was therefore investigated in two monoclonal lines of *ATM* knockout (KO) human C20 microglia-like cells^24,36^. Loss of ATM function was confirmed by the lack of phosphorylation of the ATM substrate CHK2 T68 in response to camptothecin (CPT), a topoisomerase I inhibitor known to induce ATM activation (Supplemental Figure 1A and 1B)^37^. Increased phosphorylation of STING S366, TBK1 S172 and IRF3 S396 indicated that the STING pathway is activated in both *ATM* KO C20 lines (Figure 1A). Importantly, basal expression of *IFNB1* and the ISG *IFIT2* was induced in *ATM* KO cells, however, the induction in response to stimulation with the non-cyclic dinucleotide STING agonist, diABZI^38^, was comparable to WT cells (Figure 1B).

**Figure 1:**
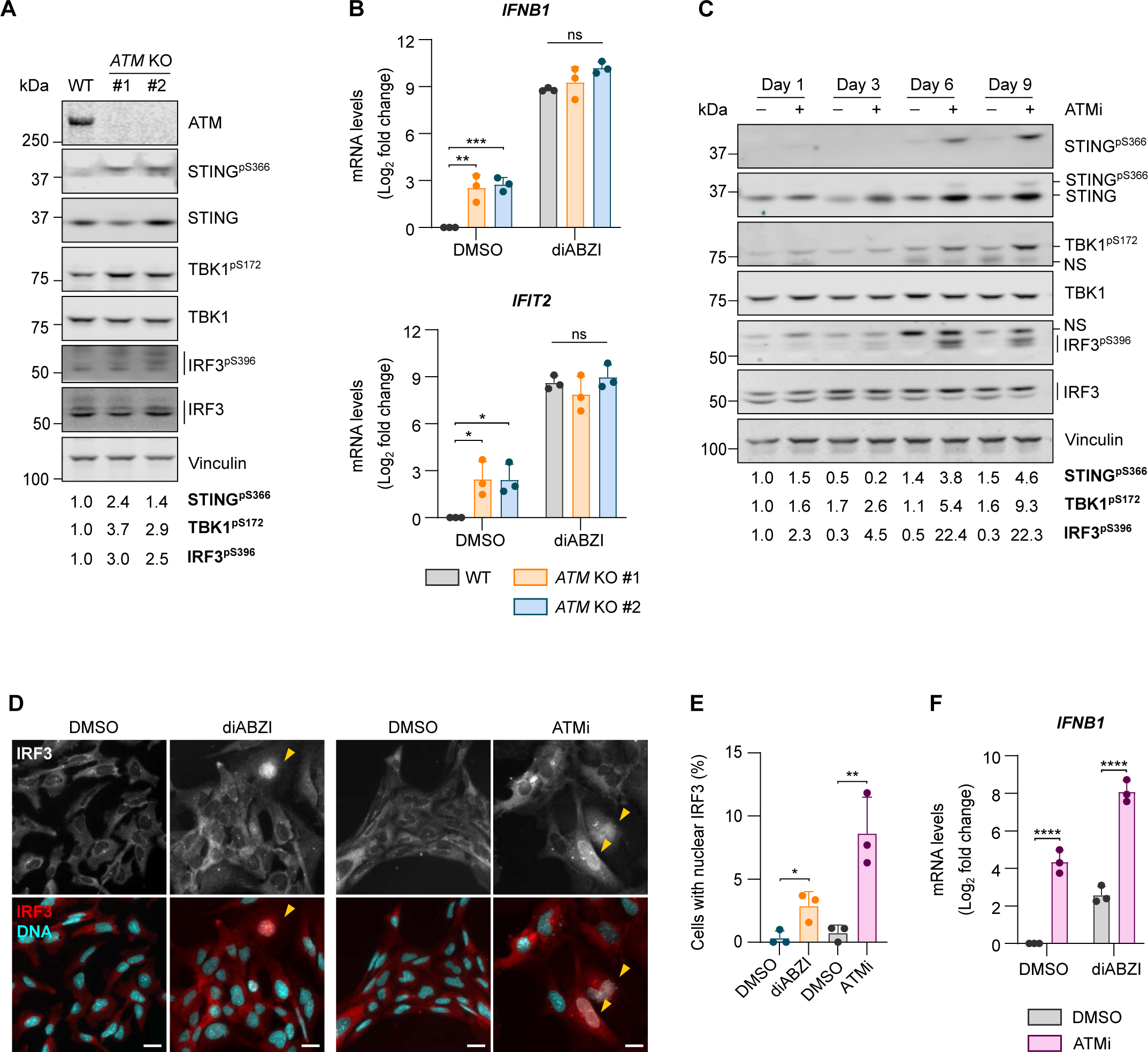
Loss of ATM function in human microglia-derived cells drives STING pathway activation and a type I interferon response. **(A)** Representative immunoblot of STING pathway activation in *ATM* KO C20 cells (clones #1 and #2). Loading control: Vinculin. Quantification of phosphorylated proteins is relative to respective total proteins and normalised to WT. **(B)** Relative mRNA levels of *IFNB1* and *IFIT2* in WT and *ATM* KO C20 cells treated with 1 µM STING agonist diABZI, or DMSO as control, for 5 h. C_q_ values normalised to *IPO8* and DMSO-treated WT cells. Mean log_2_ fold change ± SD (n=3). Two-way ANOVA with Tukey’s post-hoc comparison test. **(C)** Representative immunoblot of HMC3 cells treated with 10 nM ATM inhibitor AZD1390 (ATMi), or DMSO as control, for 1, 3, 6 or 9 days. Loading control: Vinculin. NS: nonspecific. Quantification of phosphorylated proteins is relative to respective total proteins and normalised to Day 1 DMSO control. **(D)** Representative images of IRF3 localisation in HMC3 cells stimulated for 2 h with 1 µM diABZI, or DMSO as control, or with 10 nM ATMi or DMSO for 6 days (diABZI and ATMi stimulation are part of separate experiments). Red: IRF3, cyan: DNA (DAPI). 20x magnification, scale bar = 25 µm. Yellow arrows indicate cells with nuclear IRF3. **(E)** Percentage of cells with nuclear translocation of IRF3 treated as in (D). Mean ± SD (n=3). Unpaired, two-tailed Student’s *t*-tests. **(F)** Relative *IFNB1* mRNA levels in HMC3 cells treated with ATMi or DMSO for 6 days followed by 1 µM diABZI, or DMSO as a control, for 5 h. C_q_ values normalised to *RPS13* and DMSO-treated cells. Mean log_2_ fold change ± SD (n=3). Two-way ANOVA with Tukey’s post-hoc comparison test. *p ≤ 0.05; **p ≤ 0.01; ***p ≤ 0.001; ****p ≤ 0.0001; ns = not significant.

To confirm whether loss of kinase activity induces phenotypes similar to loss of ATM protein, human microglia-derived HMC3 cells were treated with a selective ATM kinase inhibitor AZD1390 (ATMi)^39^. ATMi treatment blocked ATM auto-phosphorylation at S1981 and phosphorylation of CHK2 T68 in response to CPT, confirming successful kinase inhibition (Supplemental Figure 1C and 1D). ATMi also progressively increased levels of the DSB marker γH2AX, indicating accumulation of DNA damage (Supplemental Figure 1E and 1F). ATM inhibitor treatment led to a gradual increase in phosphorylation of STING, TBK1 and IRF3 between 3-9 days of treatment, and a concomitant upregulation of STING protein which at day 6, could be attributed to increased *STING1* transcription (Figure 1C and Supplemental Figure 1G). ATMi also induced nuclear translocation of IRF3 in a fraction of cells similar to that seen in diABZI-treated cells (Figure 1D and 1E). Low percentage of cells with nuclear IRF3 likely reflects that it can be difficult to detect owing to transient relocalisation^40^. Furthermore, ATM inhibition resulted in increased expression of the type I interferon, *IFNB1*, which was exacerbated upon stimulation with diABZI in ATMi-treated compared to control-treated HMC3 cells (Figure 1F). Different responses of ATMi-treated and *ATM* KO cells to additional stimulation with diABZI may reflect adaptation mechanisms following constitutive loss of ATM, compared to the short-term loss of function upon kinase inhibition. Taken together, these data confirm that loss of ATM function leads to activation of STING and a type I interferon response in human microglia-like cells.

### Loss of ATM function leads to cGAS accumulation at micronuclei

Micronuclei are a hallmark of genome instability known to induce cGAS-STING activation in proliferating cells^9,10^. Although microglial turnover is low in the healthy CNS, microglial proliferation can be stimulated in response to perturbations in their microenvironment. For example, rapid expansion of the population of microglia is observed following their depletion with CSF1R inhibitors in mice and in mouse models of neurodegenerative prion disease^41,42^. Moreover, cell cycle genes are upregulated in cerebellar microglia in A-T, suggesting an increase in proliferative capacity^31^.

To confirm that loss of ATM function induces the formation of micronuclei, human microglia-like HMC3 cells were treated with ATMi for 6 days followed by drug washout for 4 days (Figure 2A). As expected, loss of ATM activity led to a significant increase in micronucleated cells, which was rescued by inhibitor washout (Figure 2B and 2C). Similarly, *ATM* KO C20 lines displayed an increased micronuclei frequency compared to wild-type (WT) cells (Figure 2D and 2E). This increase was less marked than that seen with kinase inhibition, likely due to the exacerbated genome instability caused by loss of kinase activity relative to loss of ATM protein^43,44^.

**Figure 2:**
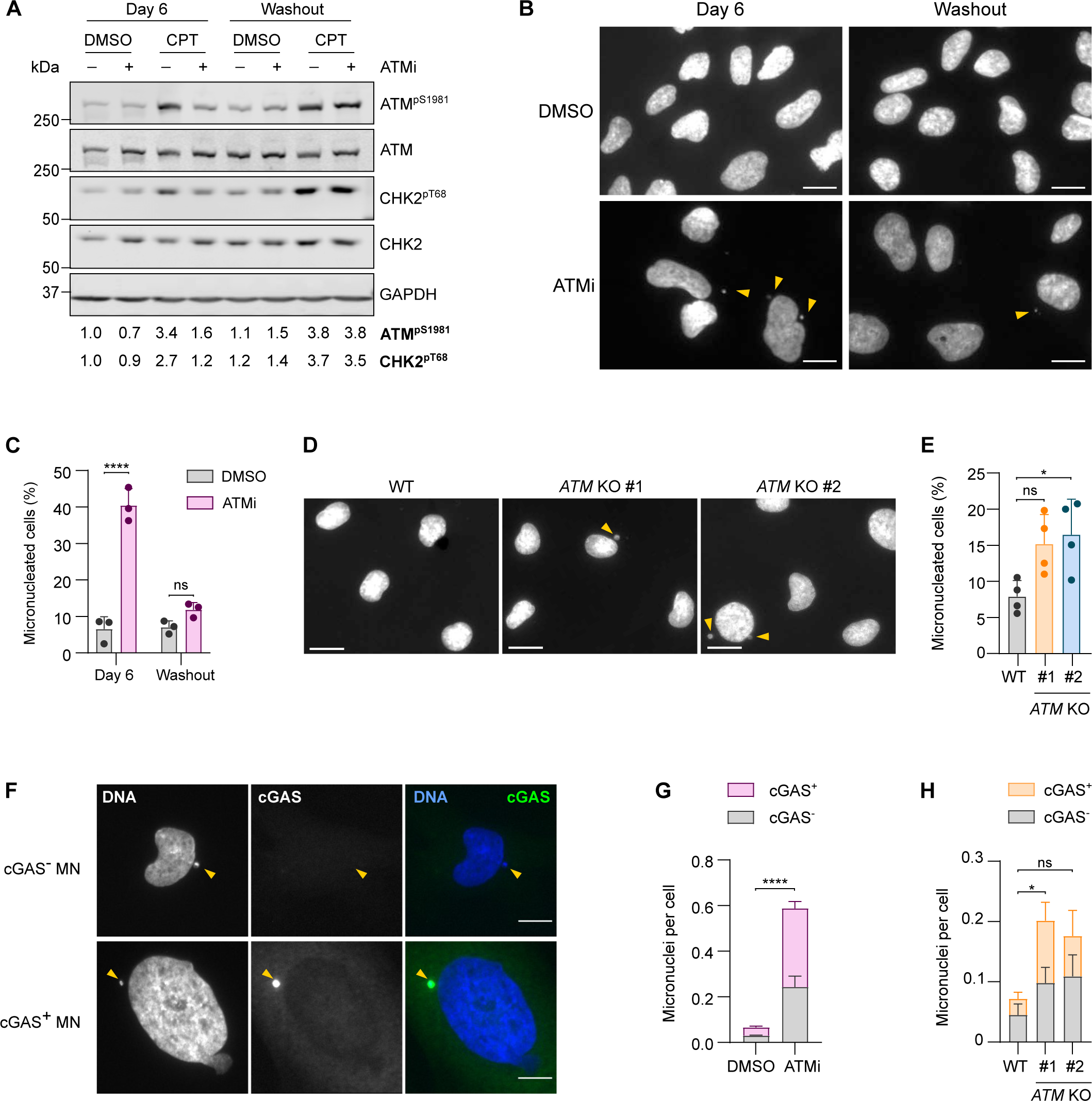
Loss of ATM function leads to cGAS accumulation at micronuclei. **(A)**Representative immunoblot of HMC3 cells treated with 10 nM AZD1390 (ATMi), or DMSO as a control, for 6 days followed by 4-day ATMi washout. To confirm ATM inhibition, cells were treated with 1 µM camptothecin (CPT) or DMSO as a control for 1 h. Loading control: GAPDH. Quantification of ATM^pS1981^ and CHK2^pT68^ is relative to respective total proteins. **(B)** Representative images of micronuclei (MN) in HMC3 cells treated as in (A). Yellow arrows indicate MN visualised by DAPI staining. 20x magnification, scale bar = 15 µm. **(C)** Quantification of MN frequency as in (B). Mean ± SD (n=3). Two-way ANOVA with Tukey’s post-hoc comparison test. **(D)** Representative images of MN in WT and *ATM* KO C20 cells. Yellow arrows indicate MN visualised by DAPI staining. 20x magnification, scale bar = 15 µm. **(E)** Quantification of MN frequency as in (D). Mean ± SD (n=4). One-way ANOVA with Tukey’s post-hoc comparison test. **(F)** Representative images of MN negative (cGAS^-^ MN) or positive (cGAS^+^ MN) for cGAS. Yellow arrows indicate MN. Blue: DAPI, green: cGAS. 20x magnification, scale bar = 15 µm. **(G)** Quantification of cGAS^+^ and cGAS^-^ MN in HMC3 cells treated with ATMi or DMSO for 6 days. MN frequency per cell is shown. Mean ± SD (n=3). Two-way ANOVA with Tukey’s post-hoc comparison test. **(H)** Quantification of cGAS^+^ and cGAS^-^ MN in WT and *ATM* KO C20 cells. MN frequency per cell is shown. Mean ± SD (n=3). Two-way ANOVA with Tukey’s post-hoc comparison test. *p ≤ 0.05; ****p ≤ 0.0001; ns = not significant.

Since cGAS can accumulate at micronuclei following micronuclear envelope collapse^9,10^, its localisation was determined in ATMi-treated HMC3 cells by immunofluorescence using a cGAS-specific antibody (Figure 2F and Supplemental Figure 2A). The number of cGAS-positive micronuclei significantly increased upon ATMi treatment as compared to control (Figure 2G). An increase in the number of cGAS-positive micronuclei was also observed in *ATM* KO C20 lines, confirming that cGAS accumulates at micronuclei in ATM-deficient human microglia-like cells (Figure 2H). However, ATM inhibition or deletion did not affect the proportion of cGAS-positive micronuclei, indicating that loss of ATM function does not alter the propensity for micronuclear envelope collapse or cGAS recruitment (Supplemental Figure 2B and 2C). Collectively, these data confirm that micronuclei are a source of cytosolic DNA in ATM-deficient microglia-like cells.

ATM loss reportedly induces accumulation of cytosolic DNA fragments and leakage of mtDNA in some cell types^6,26,33,35^. As cGAS senses dsDNA^45^, dsDNA localisation was visualised in ATMi-treated HMC3 cells by immunofluorescence confocal microscopy. Co-staining for the mitochondrial marker TOM20 confirmed that cytosolic dsDNA did not localise to the mitochondria, and loss of dsDNA signal following DNase I treatment confirmed antibody specificity. However, no increase in cytosolic dsDNA was observed upon ATM inhibition (Supplemental Figure 2D and 2E). One reported mechanism of mtDNA leakage in ATM-deficient murine cancer cells is via downregulation of the mitochondrial transcription factor A (TFAM)^33^. However, *TFAM* mRNA levels were unchanged in ATMi-treated HMC3 cells, in line with reports in Atm-deficient mouse thymocytes (Supplemental Figure 2F)^46^. These results indicate that cGAS-STING activation likely originates primarily from micronuclei formation.

To address the requirement for cell cycle progression in activating STING, mitotic entry was blocked using the CDK1 inhibitor (CDK1i) RO-3306 in the absence or presence of ATMi. Cells were treated with ATMi for 6 days, with addition of CDK1i for the final 48 h (Supplemental Figure 2G). Immunoblotting and cell cycle analysis confirmed effective ATM inhibition and synchronisation of HMC3 cells in G2 phase upon CDK1i treatment (Supplemental Figure 2H and 2I). ATM-inhibitor induced phosphorylation of STING and IRF3 and increased expression of the interferon-responsive transcription factor STAT1 were alleviated by CDK1i treatment, confirming a requirement for mitotic progression (Supplemental Figure 2J). Notably, TBK1 protein levels are mildly downregulated in synchronised cells which may partially contribute to pathway inhibition. This downregulation may be linked with previously reported roles for TBK1 in the G2/M checkpoint and microtubule dynamics during mitosis^47,48^.

Taken together, these data indicate that loss of ATM function in proliferating microglia leads to recognition of micronuclei by cGAS, providing a source of sterile STING-dependent inflammation.

### cGAS dependency of STING activation

STING activation can occur through mechanisms independent of cGAS and cytosolic DNA, such as inappropriate trafficking to the Golgi or via nuclear signalling upon sensing of DNA damage by PARP1^49,50^. Furthermore, ATM inhibition was reported to activate TBK1 in a cGAS-STING-independent manner via SRC kinase in pancreatic cancer cells^34^.

To confirm whether interferon signalling is activated via canonical cytosolic DNA sensing, cGAS and STING were depleted using siRNA in ATMi-treated HMC3 microglia. Knockdown (KD) of cGAS abolished phosphorylation of STING following ATMi treatment, while KD of cGAS or STING abolished TBK1 phosphorylation (Figure 3A and Supplemental Figure 3A). Biochemical fractionation revealed that nuclear translocation of phosphorylated IRF3 and STAT1 in ATMi-treated cells was abolished upon cGAS or STING KD (Figure 3B and Supplemental Figure 3B). Importantly, an increase in expression of *IFNB1* and *IFIT2* in ATMi-treated cells is abolished upon cGAS or STING KD, while induction of *IL6* is only partially dependent on cGAS and STING (Figure 3C). As *IL6* is a target of NF-κB transcription factors, and we previously reported activation of non-canonical NF-κB upon loss of ATM function, residual inflammatory signalling upon cGAS-STING deletion may be driven by this pathway^24^.

**Figure 3:**
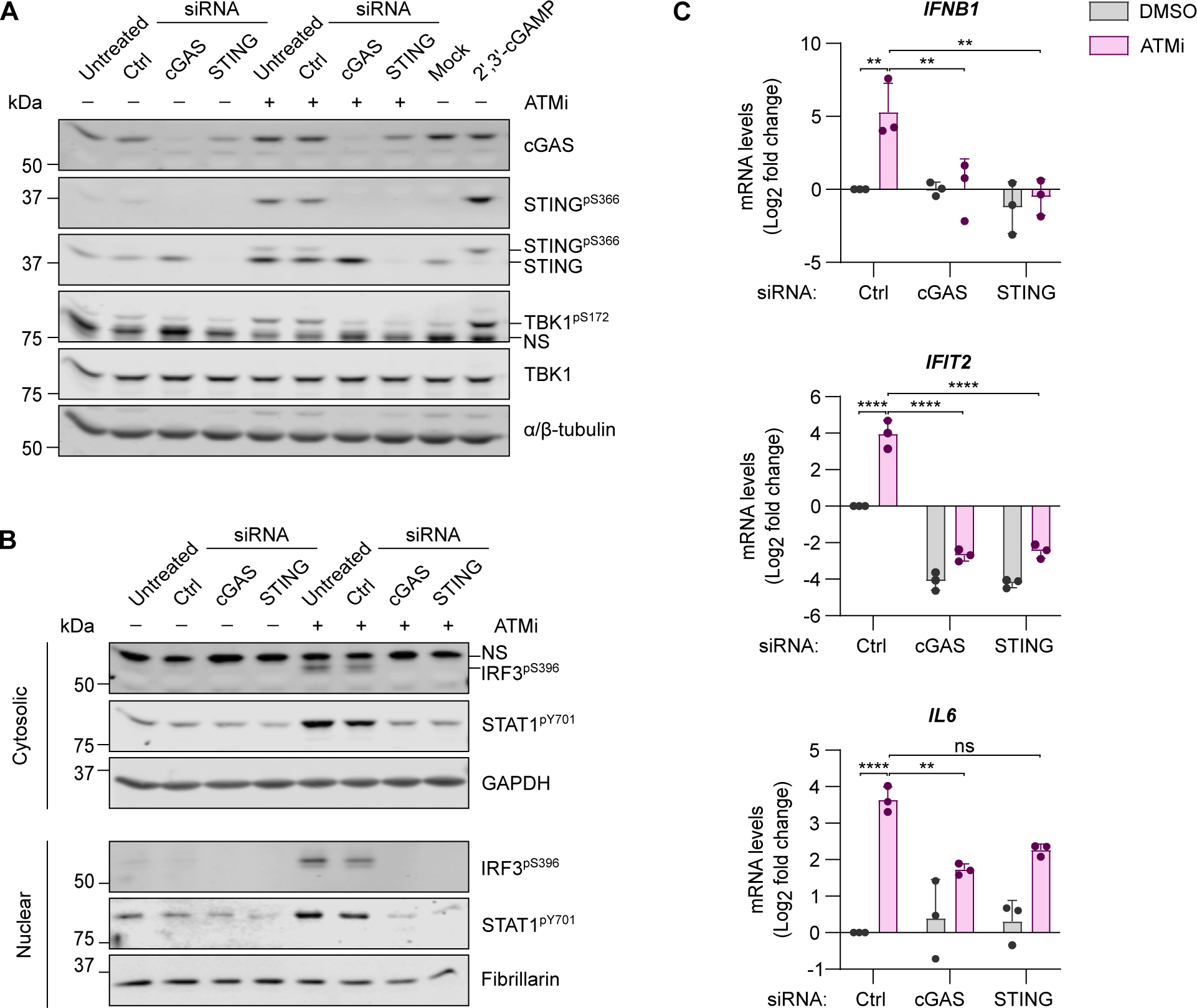
STING activation and type I interferon signalling in ATM deficiency are dependent on cGAS. **(A)**Representative immunoblot of siRNA-mediated cGAS and STING depletion in HMC3 cells treated with 10 nM ATMi, or DMSO as a control, for 6 days. Positive control: 7 µM 2’,3’-cGAMP for 1.5 h delivered by digitonin permeabilization, mock: digitonin permeabilization. Loading control: α/β-tubulin. NS: nonspecific. **(B)** Representative immunoblot of IRF3^pS396^ and STAT1^pY701^ levels in cytosolic and nuclear extracts of HMC3 cells as in (A). Cytoplasmic marker: GAPDH; nuclear marker: fibrillarin. **(C)** Relative mRNA levels of *IFNB1*, *IFIT2*, and *IL6* of cells as in (A). C_q_ values normalised to *RPS13* of cells treated with control (Ctrl) siRNA and DMSO. Mean log_2_ fold change ± SD (n=3). Two-way ANOVA with Tukey’s post-hoc comparison test. **p ≤ 0.01; ****p≤ 0.0001; ns = not significant.

Taken together, these data demonstrate that canonical cGAS-STING signalling drives the type I interferon response in ATM-deficient microglia-derived cells.

### Type I interferon signalling elicits chemokine and motility transcriptional programmes in human iPSC-derived microglia

STING signalling has been linked with activation of NF-κB signalling and induction of senescence programmes in Atm-deficient murine microglia and A-T brain organoids^26,32^. However, type I interferon is a major arm of transcriptional responses downstream of mammalian STING^51^. To determine how type I interferon shapes microglial function, bulk RNA-sequencing (RNAseq) was performed in human iPSC-derived iCell microglia treated with human IFN-β for 6 or 24 h (Figure 4A). As expected, differentially expressed genes (DEGs) in IFN-β-treated iPSC-derived microglia are enriched in gene ontology (GO) biological processes associated with antiviral immunity, including positive regulation of immune response, response to virus and positive regulation of inflammatory response (Figure 4B). Importantly, GO terms associated with cell locomotion, motility and migration were among the top 5 enriched processes. Processes linked with actin filament organisation, cell polarity and cell shape also featured in the top 20 enriched GO processes.

**Figure 4:**
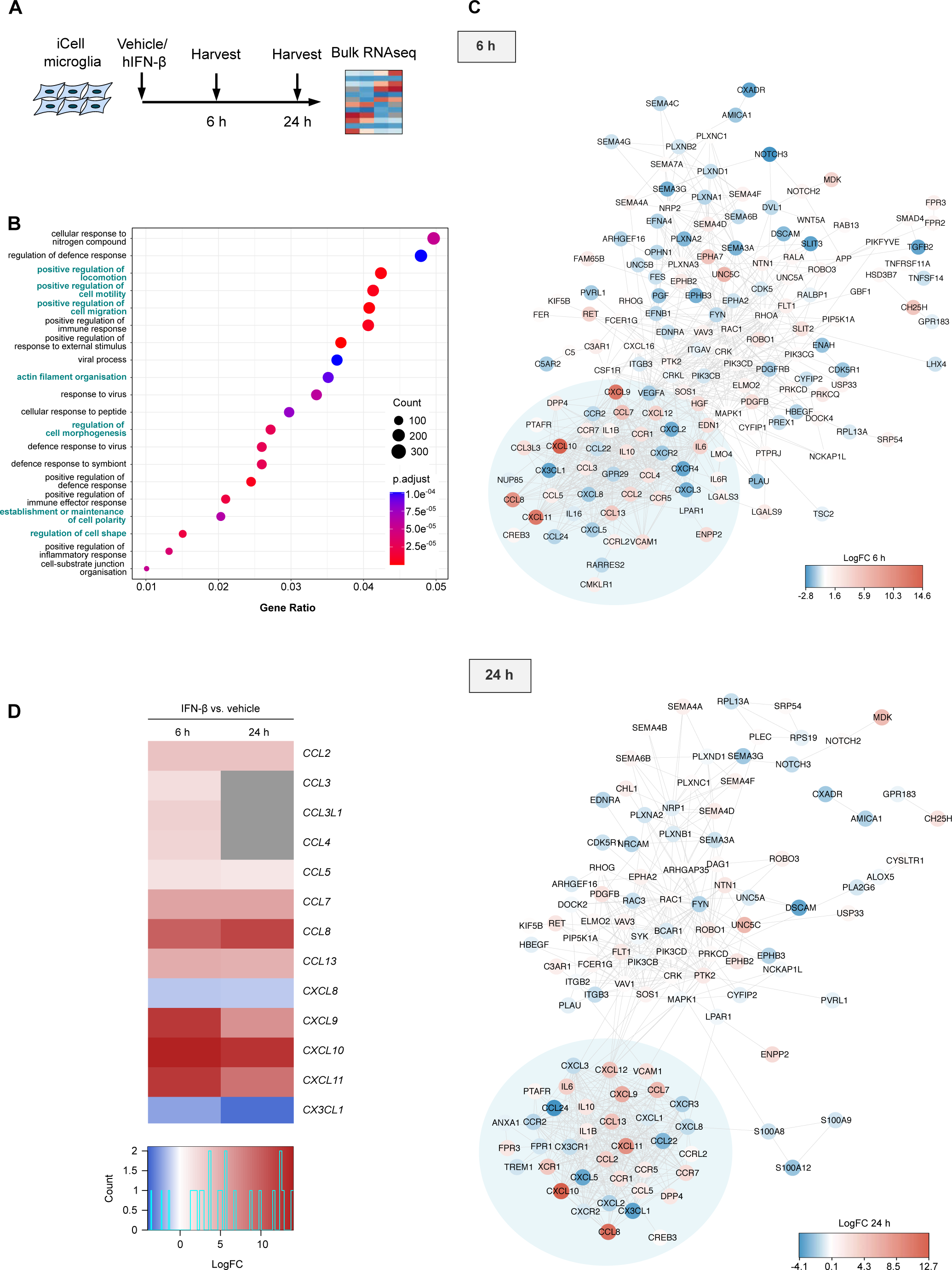
Migration-related pathways and chemotaxis genes are enriched in iPSC-derived microglia stimulated with IFN-β. **(A)** Schematic of experimental workflow. iCell microglia were treated with vehicle or 5 ng/mL human IFN-β (hIFN-β) for 6 or 24 h (n = 3), followed by bulk RNAseq. **(B)** Dotplot of top 20 GO biological processes enriched among differentially expressed genes (DEGs) in IFN-β-treated iCell microglia. Migration-related processes highlighted in teal. **(C)** STING network analysis of significantly differentially expressed genes (adjusted p-value < 0.05) related to cell chemotaxis at 6 and 24 h of IFN-β-treated compared to vehicle-treated cells. Colour scale represents log_2_ fold change (Log2FC). **(D)** Heatmap shows gene expression changes (Log2FC) of significantly differentially expressed chemokine family genes in cells treated as in (A). Grey indicates the genes that are not significantly differentially expressed (adjusted p-value > 0.05).

The chemokine superfamily is divided into C, CC, CXC, and CX3C subtypes based on the amino acid arrangement between conserved N-terminal cysteine residues. Chemokines signal via transmembrane G-protein coupled receptors (GPCRs) to regulate cell polarity, cytoskeletal organisation and adhesion, driving cell recruitment along a chemotactic gradient^52^. Protein interaction network analysis at 6 and 24 h identified a significant cluster of chemokines and their receptors, indicating enrichment of microglial chemokine signalling following IFN-β treatment (Figure 4C). Specifically, *CXCL9*, *CXCL10* and *CXCL11,* ligands of the CXCR3 receptor, and *CCL8*, a ligand of CCR2, are among the top upregulated chemokines upon IFN-β treatment at both 6 and 24 h (Figure 4D). In addition, *CCL5* among other chemokines of the CC family, is highly upregulated (Figure 4C and 4D). Therefore, chemokine signalling is a critical outcome of type I interferon responses in human microglial cells.

### ATM loss drives STING-dependent chemokine production in microglia-derived cells

Given the induction of chemokine and cell motility programmes in response to IFN-β treatment in iPSC-derived microglia, we sought to investigate whether ATM loss induces microglial chemokine production and whether this could be STING dependent. Single-cell RNAseq previously identified *Cxcl10* and *Ccl5* as signature ISGs induced specifically in micronucleated cells^10^, therefore mRNA levels of these chemokines were assessed in ATM-deficient microglia-like cells. *CCL5* and *CXCL10* were found to be significantly upregulated in *ATM* KO C20 cells and induced by STING activation with diABZI (Figure 5A). As siRNA delivery and lipofection can activate the innate immune system^53,54^, *STING1* KO HMC3 cells were established to further investigate the consequences of chronic STING activation on microglial function. In two monoclonal *STING1* KO lines, TBK1 S172 and IRF3 S396 phosphorylation was abolished (Supplemental Figure 4A and 4B) and perinuclear STING clusters were no longer detectable in response to diABZI, confirming the lack of functional STING (Supplemental Figure 4C).

**Figure 5:**
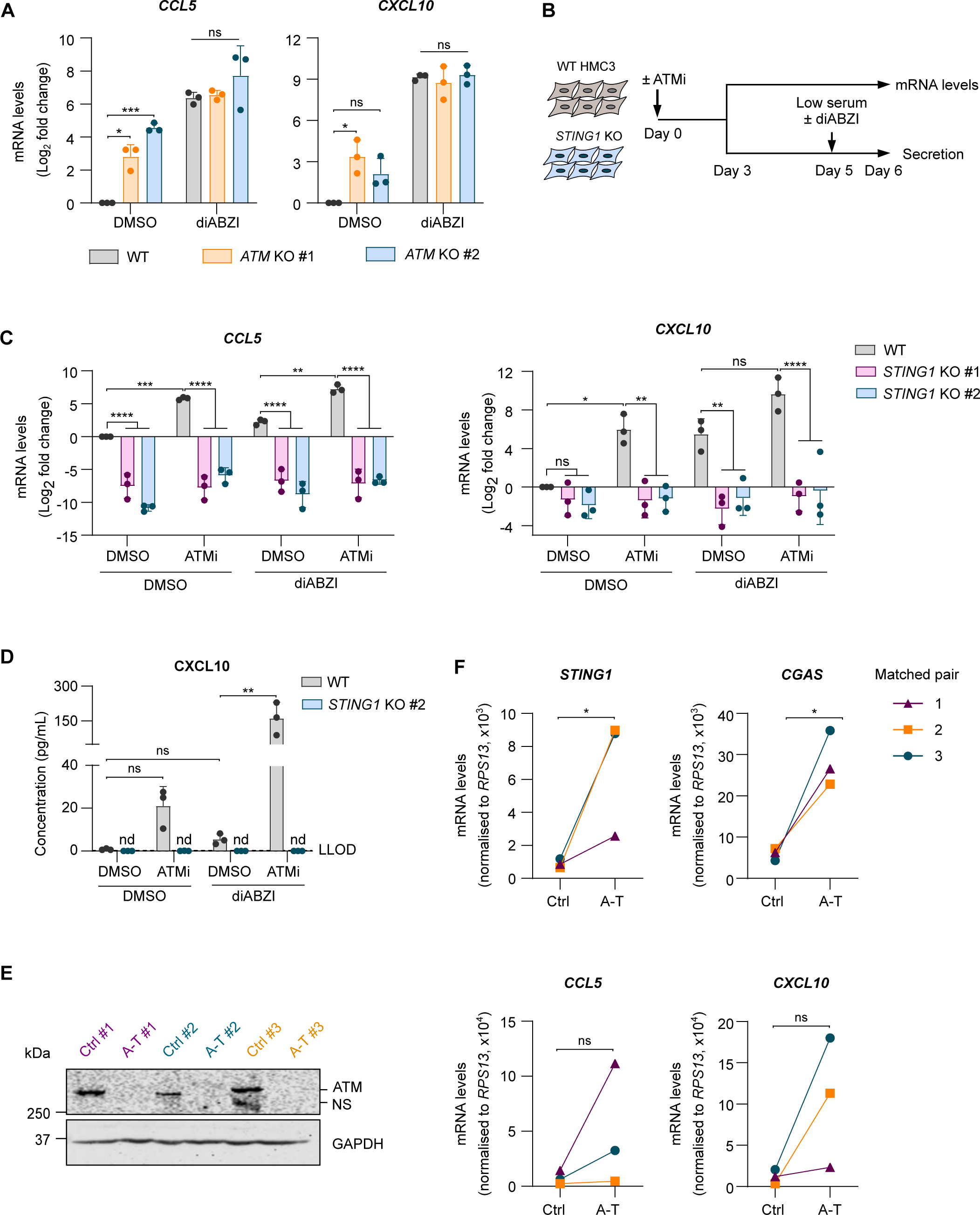
Loss of ATM function leads to STING-dependent chemokine secretion. **(A)**Relative mRNA levels of *CCL5* and *CXCL10* in WT and *ATM* KO C20 cells and upon treatment with 1 µM diABZI, or DMSO as control, for 5 h. C_q_ values normalised to *IPO8* and DMSO-treated WT cells. Mean log_2_ fold change ± SD (n=3). Two-way ANOVA with Tukey’s post-hoc comparison test. **(B)** Schematic of experimental workflow. WT and *STING1* KO HMC3 cells were treated with 10 nM ATMi, or DMSO as a control, for 6 days. mRNA levels were measured by RT-qPCR. Secretion was measured 24 h after media change to low FBS supplemented with 1 µM diABZI, or DMSO as a control. **(C)** Relative mRNA levels of *CCL5* and *CXCL10* from cells treated as in (B). C_q_ values normalised to *RPS13* and DMSO-treated WT cells. Mean log_2_ fold change ± SD (n=3). Two-way ANOVA with Tukey’s post-hoc comparison test. **(D)** CXCL10 concentration in cell culture supernatants. Mean ± SD (n=3). Dashed line indicates lower limit of detection (LLOD) = 0.12 pg/mL. Nd: not detected. One-way ANOVA with Tukey’s post-hoc comparison test in WT cells. **(E)** Representative immunoblot of cerebellar homogenates from individuals with A-T and matched individuals with no known diseases of the CNS as controls (Ctrl). Loading control: GAPDH. NS = nonspecific. **(F)** Relative mRNA levels of *STING1*, *CGAS*, *CCL5,* and *CXCL10* as in (E). C_q_ values normalised to *RPS13*. Mean 2^-δCT^ ± SD shown (n=3). Paired *t*-test. *p ≤ 0.05; **p ≤ 0.01; ***p ≤ 0.001; ****p ≤ 0.0001; ns = not significant.

We therefore focused on the role of STING in chemokine production in basal conditions and upon treatment of WT and *STING1* KO HMC3 cells with ATMi (Figure 5B). As expected, the induction of *IFNB1* in cells treated with ATMi or diABZI was completely abolished in *STING1* KO lines (Supplemental Figure 4D). Although *IFNB1* mRNA was induced, ATM inhibition increased secretion of IFN-α2a rather than IFN-β, and this was dependent on STING (Supplemental Figure 4E). This discordance between mRNA levels is likely to be attributed to the kinetics of the secretory response. Importantly, ATM inhibition also induced a robust chemokine response, evidenced by increased mRNA levels of *CCL5* and *CXCL10*, which were fully dependent on STING. Stimulation with diABZI similarly induced *CCL5* and *CXCL10* expression (Figure 5C). Moreover, *STING* deletion markedly reduced *CCL5* mRNA in basal conditions, and a similar tendency was observed in the case of *CXCL10*, suggesting tonic, i.e. basal, STING-dependent chemokine signalling. As *CXCL10* was the most strongly upregulated gene following IFN-β treatment in iPSC-derived microglia, its concentrations were measured in cell culture supernatants. ATM inhibition promoted CXCL10 secretion in a STING-dependent manner, whereas CXCL10 was not detected in supernatants from *STING1* KO cells (Figure 5D).

Given the robust chemokine response *in vitro*, mRNA levels of *CGAS*, *STING1*, *CCL5* and *CXCL10* were determined by RT-qPCR in cerebellar homogenates from individuals with A-T and from age- and gender-matched individuals with no known diseases of the CNS (Figure 5E and 5F). Confirming previous single-nucleus RNAseq data, expression of *CGAS* and *STING1* was significantly upregulated in A-T cerebella compared to matched controls^31^. Importantly, levels of *CCL5* and *CXCL10* were also increased in A-T brains, albeit not significantly, likely owing to genetic variability between individuals and a small sample size limited by tissue availability (Figure 5F). Whole cerebellum gene expression changes therefore indicate upregulation of *CGAS, STING1,* and possibly chemokines in A-T. Taken together, these data demonstrate that STING drives production of chemokines upon loss of ATM function in human microglia-like cells with potential relevance in human disease.

### Persistent DNA damage promotes a STING-dependent chemokine response

To investigate the role of DNA damage in STING-dependent chemokine responses, WT and *STING1* KO HMC3 cells were treated with the topoisomerase II poison etoposide, the DNA synthesis inhibitor aphidicolin and the DNA crosslinking agent mitomycin C. To model persistent low-level DNA damage, cells were treated for 6 days using low doses of drugs that did not induce cell death (Supplemental Figure 5A). All DNA damaging agents induced the formation of micronuclei and consequent STING activation, as evidenced by increased STING S366 and TBK1 S172 phosphorylation, which was absent in *STING1* KO cells (Supplemental Figure 5B and 5C). Moreover, all three agents induced *CCL5* expression, which was dependent on STING (Supplemental 5D). These data indicate that DNA damage, rather than ATM loss per se, drives STING-dependent chemokine induction in human microglia-derived cells.

### ATM inhibition drives microglial chemotaxis in a STING-dependent manner

Microglia are highly motile cells, as their processes dynamically survey their microenvironment in the absence of a stimulus, and are rapidly directed towards sites of infection or injury^55^. Given that STING regulates tonic chemokine production in microglia-like cells, the role of STING in regulating basal microglial migration was investigated. To address this, a variant of the scratch-wound assay was utilised, in which cells are seeded in a well containing silicone inserts which are subsequently removed to produce a uniform cell-free gap (Figure 6A). Live cell imaging showed that deletion of *STING1* leads to delayed gap closure, indicating that STING regulates basal microglial migration (Figure 6B and 6C). Furthermore, chemotactic migration of WT and *STING1* KO HMC3 cells, which were grown in the presence or absence of ATM inhibitor for 6 days, was investigated using xCELLigence label-free real-time cell analysis based on measuring cellular impedance (Figure 6D)^56^. Loss of ATM activity was demonstrated to enhance chemotactic migration of cells towards their respective conditioned medium, and this effect was reduced in the absence of STING (Figure 6E).

**Figure 6:**
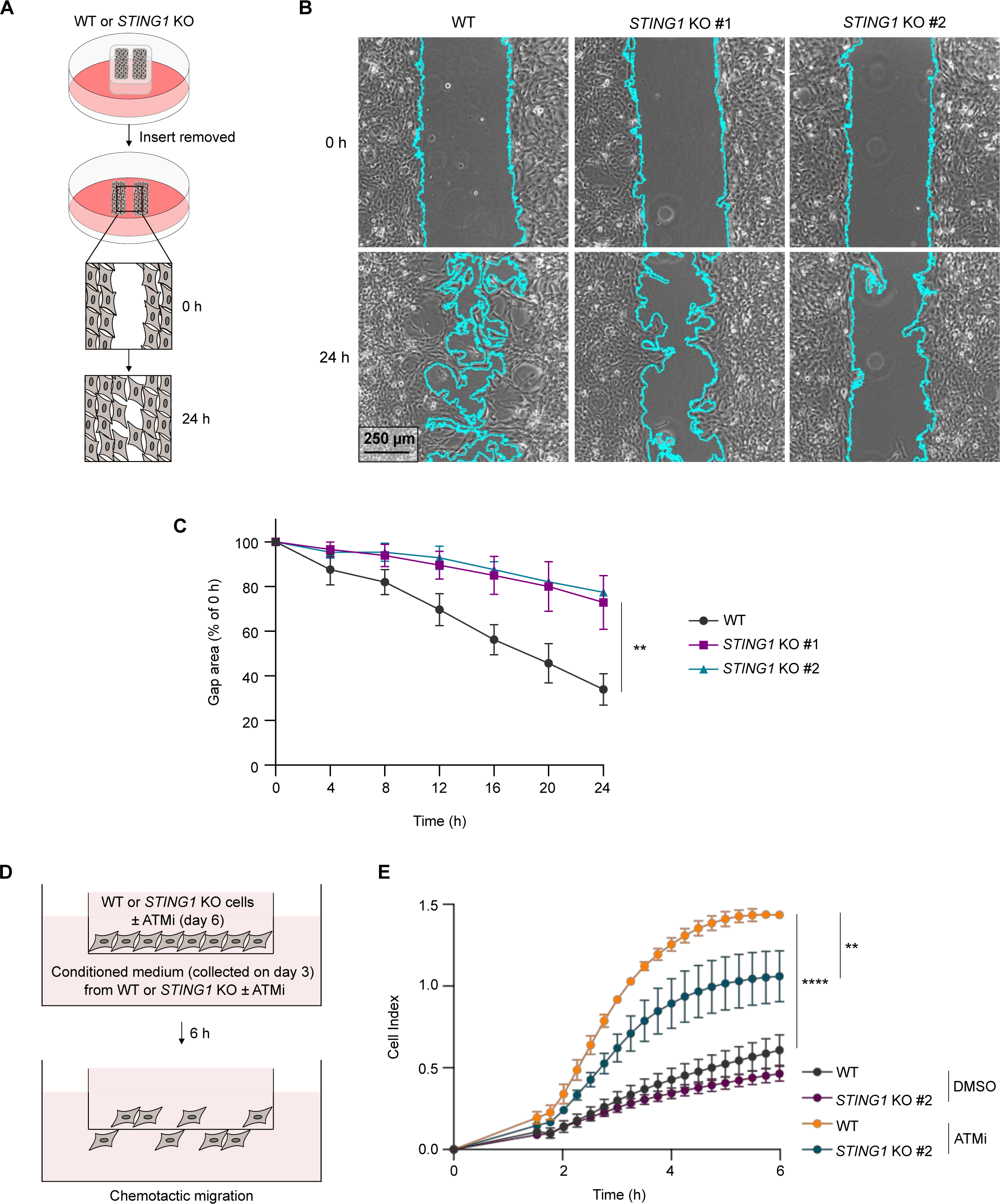
STING regulates microglial chemotaxis. **(A)**Schematic of experimental workflow for wound healing migration assay. **(B)** Representative phase contrast microscopy images of gap area at 0 and 24 h in WT and *STING1* KO HMC3 cells. 4x magnification, scale bar = 250 µm. **(C)** Normalised gap area expressed as a percentage of gap area at 0 h. Mean ± SD (n = 3). Area under curve analysed using one-way ANOVA with Tukey’s post-hoc comparison test. **(D)** Schematic of experimental workflow for xCELLigence chemotaxis assay. WT and *STING1* KO HMC3 cells were treated with 10 nM ATMi, or DMSO as a control, for 6 days. Chemotactic migration towards the respective conditioned medium was measured. **(E)** Kinetics of microglial chemotaxis of cells as in (D). Mean ± SD (n = 3 technical replicates). Representative of 3 independent experiments. Two-way ANOVA with Tukey’s post-hoc comparison test. **p ≤ 0.01; ****p ≤ 0.0001.

Overall, these data demonstrate that STING regulates chemotaxis of microglia-derived cells in basal conditions and following loss of ATM function.

## DISCUSSION

In this study, we utilised A-T as a model genome instability disorder to explore how cGAS-STING signalling regulates microglial function upon persistent DNA damage. In line with previous observations upon ATM deletion or pharmacological inhibition, we find that the cGAS-STING pathway induces chronic inflammation following ATM loss in models of human microglia-derived cells^6,26,32,35^. Both genomic and mitochondrial origins of cytosolic DNA in ATM deficiency have previously been proposed^12,33,35^. Whilst a mitochondrial origin cannot be ruled out in the present work, we find that STING signalling in the absence of ATM is associated with cGAS activation by micronuclei in proliferating microglia (Figure 7). Indeed, ATM-deficient cells present with chromosomal rearrangements that form upon non-homologous end joining of single-ended DSBs and could drive micronucleus formation^57^. Evidently, STING-dependent inflammation is phenocopied upon prolonged treatment of ATM-proficient microglia-derived cells with a range of DNA damaging agents, all of which could potentiate the formation of DSBs and consequently micronuclei. These observations argue against a specific effect of ATM loss as a driver of STING activation in microglia. We therefore propose that chronic inflammation in ATM deficiency is driven by genome instability, in line with the nuclear origin of cytosolic DNA upon Atm inhibition in murine microglia^12^.

**Figure 7:**
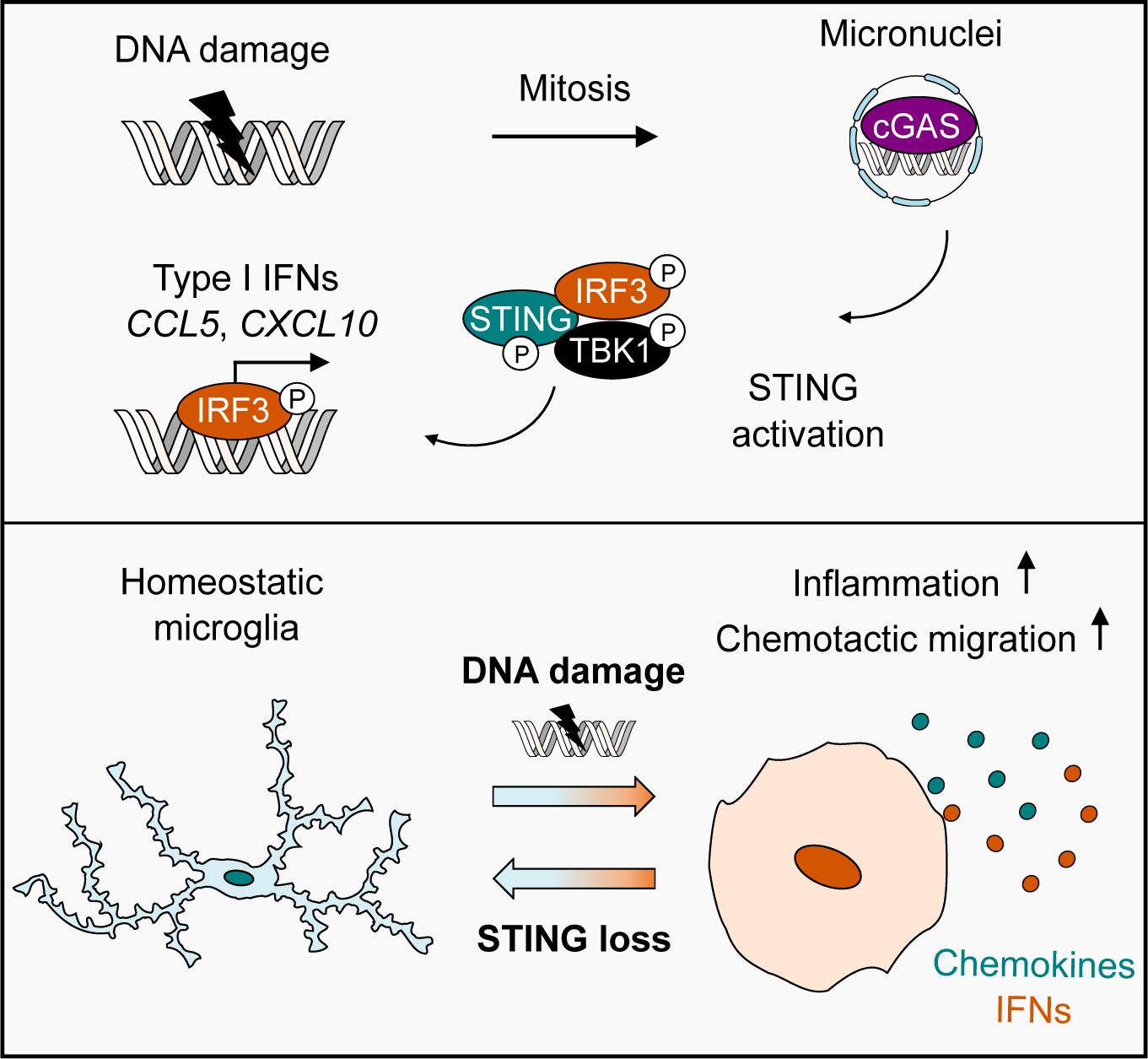
Proposed model of STING-dependent regulation of microglial function in genome instability. Persistent DNA damage, such as that observed in ATM deficiency, leads to formation of micronuclei in proliferating microglia. Activation of the cGAS-STING pathway culminates in IRF3 activation and production of type I interferons and chemokines, including *CCL5* and *CXCL10*. Secreted inflammatory mediators likely mediate a transition from homeostatic functions of microglia to chronic inflammation and chemotaxis to reactive microglia via autocrine and/or paracrine signalling, promoting chronic inflammation and chemotaxis. Blockade of STING rescues this process and has potential to alleviate perturbations to CNS homeostasis in genome instability.

As micronuclei form during mitosis, the cellular proliferative capacity will dictate the rate of micronuclei formation. Importantly, microglial proliferation is highly responsive to external cues, including inflammation, and expansion of the microglial compartment is frequently observed in neurodegenerative disease^41,58^. It is therefore possible that microglial proliferation is stimulated by perturbations in the CNS microenvironment during genome instability, perpetuating neuroinflammatory processes. Consistent with this, microgliosis is observed in the spinal cord of Atm-deficient rats and several studies have highlighted increased turnover of cerebellar microglia relative to cortical microglia, suggesting a region-specific increase in microglial proliferative capacity^25,30,31^.

Previous studies established a connection between genome instability, activation of STING, and inflammation in ATM deficiency^12,26,32,35^. We extend these findings by demonstrating that interferon responses to ATM inhibition are abolished in STING-deficient cells, however, the consequences of type I interferon production on microglial homeostasis are unclear. We therefore focus on investigating interferon-dependent regulation of microglial function. In line with IFN-α administration in murine microglia, we discover that IFN-β potently upregulates signalling networks associated with chemotaxis and migration in human iPSC-derived microglia^59,60^. These data identify chemotactic responses as a key outcome of type I interferon signalling. Indeed, STING signalling induces production of chemokines, such as *CCL5* and *CXCL10*, in human microglia-like cells upon ATM loss or persistent DNA damage (Figure 7). Whilst acute chemotactic responses are critical for antimicrobial immunity, excessive and/or chronic production of chemokines is implicated in detrimental microglial functions *in vivo*. For example, CXCL10-CXCR3 signalling regulates microglial recruitment and dendritic loss following axonal injury^61^. Moreover, microglia-derived CCL3/4/5 activates neuronal CCR5, leading to autophagy inhibition and accumulation of protein aggregates in mouse models of Huntington’s disease and tauopathy^62^. Interestingly, CCR5 also promotes microglial recruitment to cerebral vasculature to support the blood brain barrier (BBB) during acute inflammation, however, chronic inflammation drives BBB dysfunction via microglial phagocytosis of astrocytic end feet^63^. Sustained, excessive chemokine production may therefore shift the balance towards eliciting deleterious effects on neuronal function and the CNS microenvironment. As chemokines are implicated in recruitment of peripheral immune cells, the question of whether cGAS-STING signalling drives immune infiltration during chronic CNS inflammation in A-T warrants further investigation^64^.

Importantly, this study identifies STING as a regulator of microglial chemotaxis, both in basal conditions and during ATM deficiency (Figure 7). In mice, STING activation mediates chemotactic recruitment of T-cells into the inflamed peritoneum and tumour-infiltrating neutrophils^65,66^. Conversely, STING activation supresses migration of cancer cells and tumour-infiltrating neutrophils by reducing translation of the matrix-remodelling serine peptidase PLAU^67^. However, knowledge of whether STING regulates microglial migration is lacking. We find that STING-deficient microglia-like cells display a marked defect in migration in the absence of stimulation. Whilst STING may act intrinsically to regulate cytoskeletal dynamics, it is possible that tonic chemokine signalling underlies this phenotype, as STING loss reduced basal chemokine expression. Previous work identified enhanced migration in Atm-deficient primary murine microglia and astrocytes isolated from the cerebellum, but not the cerebral cortex^27^. This indicates cerebellum-specific effects of ATM loss on microglial migration may exist, and we propose that STING signalling may underlie this process. In addition, the Rho family of small GTPases coordinate actin cytoskeleton dynamics during cell migration. In ATM-deficient human fibroblasts, oxidative stress reportedly promotes increased migration by activating the Rho GTPase Rac1, suggesting that oxidative stress may also factor into regulation of cell migration upon ATM loss^68^.

Microglial soma are largely immobilised during baseline microglial motility *in vivo*, however, their processes are highly motile enabling constant surveillance of the local parenchyma^55^. Uniquely, cerebellar microglia exhibit baseline soma motility and frequently interact with Purkinje cell soma and dendrites^69^. Enhanced sampling of the surrounding parenchyma may improve scavenging of apoptotic cells or debris and detection of pathogenic stimuli, promoting beneficial outcomes in an acute setting. Conversely, increased surveillance in the degenerating brain may amplify recognition of neuronal stress, including via chemotactic-driven clustering of activated microglia, which might potentiate detrimental functions of microglia. For example, we previously showed that ATM-deficient microglia form clusters which damage neurons *in vitro* through aberrant engulfment^24^. Finally, cytoskeletal reorganisation required for sustained chemotaxis is energetically expensive and may drive metabolic reprogramming and microglial dysfunction.

In conclusion, this study identifies cytosolic DNA sensing as a critical mechanism regulating microglial motility and chemotaxis in health and disease. Given the expanding roles of cGAS-STING signalling in neurodegeneration^70^, our study uncovers mechanistic insights into inflammatory mechanisms not just in A-T, but more broadly in neurological diseases associated with genome instability and excessive interferon production in the CNS. This work therefore provides a rationale for exploring STING inhibition as a therapeutic strategy to alleviate neurological decline in A-T and similar genome instability disorders.

## MATERIALS AND METHODS

### Cell culture

Immortalised foetal human microglial HMC3 cells were kindly gifted by Dr Brian Bigger, University of Manchester^71^. HMC3 cells were maintained in DMEM (4.5 g/L glucose, GlutaMAX^TM^ supplement, no pyruvate; Gibco™) supplemented with 10% (v/v) heat-inactivated FBS (HI-FBS; Gibco). Immortalised adult human C20 microglial cells were kindly provided by Dr David Alvarez Carbonell, Case Western Reserve University^72^. C20 cells were cultured in DMEM:F-12 with 3.151 g/L glucose, L-glutamine and 15 mM HEPES (Merck) supplemented with 1% (v/v) HI-FBS and N-2 supplement (Gibco™). *ATM* knockout (KO) C20 cells generated using CRISPR-Cas9 were described previously^24^.

All cell lines were cultured at 37°C in 5% CO_2_ and 95% humidity. Cells were routinely tested for mycoplasma and experiments were performed below passage 35.

### Generation of *STING1* KO HMC3 cells

*STING1* KO HMC3 cells were generated using the pSpCas9(BB)-2A-Puro (PX459) V2.0 vector kindly gifted by Feng Zhang (Addgene plasmid #62988)^73^ containing the following single guide RNA (sgRNA) sequence targeting *STING1*: 5’-GCTGGGACTGCTGTTAAACG-3’. The vector was delivered into HMC3 cells by Magnetofection™ using Glial-Mag (Oz Biosciences) according to manufacturer’s instructions. Cells were then selected with 1.2 µg/mL puromycin for 3 days before single cell cloning.

Clones were screened by fluorescent PCR-capillary electrophoresis. DNA was extracted from clones using QuickExtract™ DNA Extraction Solution (Lucigen) according to the manufacturer’s protocol. A region flanking the sgRNA target site was amplified using a 5’-FAM labelled forward primer (5’-FAM-GCTAGGCATCAAGGGAGTGA-3’) and unlabelled reverse primer (5’-TGGATTTCTTGGTGCCCACA-3’). PCR was performed using Phusion polymerase (Invitrogen) with the following cycling parameters: denaturation (98°C, 30 s); 35 cycles of denaturation (98°C, 10 s), annealing (62°C, 30 s) and extension (72°C, 15 s); final extension (72°C, 5 min). Fragment analysis was performed by capillary electrophoresis on a 3100 XL Genetic Analyser (Life Technologies) by the Department of Biochemistry sequencing facility (University of Cambridge, UK). Successful KO of *STING1* was confirmed by Sanger sequencing and western blotting.

### RNA interference

siRNA duplexes were synthesised by Merck and used at a final concentration of 25 nM unless stated otherwise (sequences are listed in Supplemental Table 1). siRNA was delivered by forward transfection using Lipofectamine® RNAiMAX™ Transfection Reagent (Invitrogen^TM^) according to manufacturer’s instructions. Media was changed 8 h post-transfection to limit toxicity and cells were harvested 72 h post-transfection.

### Chemical treatments

The ATM inhibitor AZD1390 (ATMi; Selleck Chemicals) was used at 10 nM for 1 – 9 days with replenishment of the inhibitor every 2 days. To induce ATM activation, cells were treated with 1 µM camptothecin (CPT; Cayman Chemical Company) for 1 h. The STING agonist diABZi (Selleck Chemicals) was used at 1 µM. To induce DNA damage, cells were treated with 250 nM Etoposide (ETO; Cayman Chemical Company), 30 nM Mitomycin C (MMC; Santa Cruz), or 150 nM Aphidicolin (APH; Merck) for 6 days. To induce apoptosis, cells were treated with 1 µM staurosporine for 6 h. Treatment times are indicated in figure captions. For compounds dissolved in DMSO, DMSO concentration was equivalent between conditions and <0.1% (v/v).

For 2’,3’-cGAMP stimulation, cells were washed in PBS then incubated for 3 min at 37°C in digitonin permeabilization buffer (50 mM HEPES pH 7.0, 100 mM KCl, 3 mM MgCl_2_, 0.1 mM DTT, 85 mM sucrose, 0.2% (w/v) BSA, 1 mM ATP, 0.1 mM GTP, 4 µM digitonin) with or without 7 µM 2’,3’-cGAMP (Invivogen). Cells were further washed in PBS and incubated in culture medium until collection. Incubation times post-treatment are indicated in figure legends.

### Cell synchronisation in G2 cell cycle phase

Exponentially growing cells were pre-treated with 10 nM ATMi or DMSO as a control for 24 h, then treated for an additional 24 h in the presence (synchronised in G2 cell cycle phase) or absence (asynchronised) of 9 μM CDK1 inhibitor RO-3306 (CDK1i; Tocris Bioscience) prior to collection.

### Cell cycle analysis

Cells were pulsed with 10 µM BrdU for 30 min, then collected by trypsinisation and fixed in ice-cold 70% ethanol for at least 30 min. After removal of ethanol by centrifugation, cells were incubated in 2 M HCl containing 0.1 mg/mL pepsin for 20 min at 37°C. Cells were washed several times in PBS and 1% (v/v) HI-FBS in PBS, collected by centrifugation at 350 *g* for 5 min and incubated with anti-BrdU antibody at 1:100 dilution in 1% (v/v) HI-FBS-PBS for 1 h at 22°C. Cells were washed in PBS and incubated with AlexaFluor 488 goat anti-mouse secondary antibody diluted 1:200 in 1% (v/v) HI-FBS-PBS for 45 min at 22°C. Samples were further treated with 100 μg/mL RNase A (QIAGEN) for 30 min at 37°C, supplemented with 100 ng/mL propidium iodide (PI) and analysed on the green FL1-A and red FL2-A channels using an Accuri™ C6 Plus flow cytometer (BD Biosciences).

### Preparation of whole cell lysates

Cells were lysed in ice-cold RIPA buffer (25 mM Tris-HCl, pH 8.0, 150 mM NaCl, 1% (v/v) Triton X-100, 1% (w/v) sodium deoxycholate, 0.1% (w/v) sodium dodecyl sulphate (SDS), 5 mM EDTA) supplemented with 1 mM phenylmethylsulfonyl fluoride, 1 µM staurosporine, 1 µg/mL each of aprotinin, chymostatin, leupeptin, pepstatin, N-ethylmaleimide and 1x phosphatase inhibitor cocktail (Calbiochem). Lysates were incubated with rotation for 30 min at 4°C and centrifuged at 20,000 *g* for 20 min at 4°C. The protein concentration of supernatants was measured using Bradford assay reagent (Bio-Rad). Lysates were prepared at equal concentrations and boiled at 95°C for 10 min in denaturing loading buffer (25 mM Tris-HCl, pH 6.8, 2.5% v/v β-mercaptoethanol, 1% (w/v) SDS, 5% (v/v) glycerol, 0.05 mg/mL bromophenol blue, 1 mM EDTA) before SDS-PAGE.

### Biochemical fractionation

Cell pellets were harvested by trypsinisation and resuspended in plasma membrane lysis buffer (20 mM Tris-HCl, pH 7.4, 2.5 mM MgCl_2_, 0.3% (v/v) NP-40) supplemented with protease/phosphatase inhibitors as above. After incubation on ice for 15 min, lysates were centrifuged for 10 min at 1,500 *g*, 4°C. The supernatant served as the cytoplasmic fraction and the pellet containing nuclei was resuspended in 20 mM phosphate buffer, 0.5 M NaCl, 1 mM EDTA, 0.75% (v/v) Triton X-100, glycerol 10% (v/v) and 5 mM MgCl_2_. After incubation on ice for 15 min, nuclear lysates were centrifuged for 10 min at 20,000 *g* to obtain the nuclear fraction. The protein content of cytosolic fractions was determined by Bradford and equal volumes of cytoplasmic and nuclear fractions were analysed by SDS-PAGE.

### Immunoblotting

Proteins were separated using 10 or 4-16% Tris-glycine SDS-PAGE and transferred onto Immobilon®-FL PVDF membranes, which were blocked using Intercept®-TBS blocking buffer (LI-COR Biosciences). Primary antibodies were diluted in Intercept®-TBS, 0.1% (v/v) Tween®20 and secondary antibodies were diluted in Intercept®-TBS, 0.1% (v/v) Tween®20, 0.01% (w/v) SDS. Antibodies are detailed in Supplemental Table 1. Membranes were visualized using an Odyssey® CLx Imaging System and quantification was performed using Image Studio™ Lite Software (both LI-COR Biosciences). Band intensities were normalised to loading controls, or to the corresponding total protein for phosphorylated proteins, and presented as fold-change relative to the experimental control. Immunoblots are representative of at least 3 independent experiments.

### Brain tissue homogenisation and processing

Blocks of fresh frozen cerebellar tissues from individuals with A-T and that of individuals with no known diseases of the CNS (controls) were obtained from the NIH Neurobiobank at the University of Maryland, Baltimore, MD (United States). The use of samples was covered by the University of Maryland’s IRB approval no. HP-00042077 and the HTA site license no. 12196 that permits work with human tissue at the Department of Biochemistry (University of Cambridge, UK). The A-T and control tissue pairs were matched by age and gender (Supplemental Table 2). The tissues were cut into 20-µm thick sections with a cryostat (Leica CM 1950) maintained at −20°C. To extract the RNA, 5-6 sections were collected into a tube containing lysing matrix beads (1.4 mm ceramic spheres, MP Biologicals), supplemented with 1 mL of the TRI Reagent® (Merck), and snap frozen on dry ice. After thawing, samples were homogenized by vortexing and the RNA was extracted as previously described^74^.

To extract proteins, tissue sections were collected into tubes with lysing matrix beads (as above) containing RIPA buffer supplemented with protease/phosphatase inhibitors as above and snap frozen in liquid nitrogen. After thawing on ice, tissues were homogenized by intermittent vortexing at 4°C, then centrifuged at 20,000 *g* for 20 min at 4°C. Protein lysate concentrations were measured and prepared for SDS-PAGE analyses as above.

### RT-qPCR

Total RNA from adherent cells was extracted using the RNeasy Mini Kit (QIAGEN) and treated with DNase I (ThermoFisher) according to manufacturer’s instructions. RNA integrity was assessed by agarose gel electrophoresis and purity was determined by measuring the ratio of absorbance at 260/280 (A_260/280_) and 260/230 (A_260/230_) using a NanoDrop™ ND-1000 Spectrophotometer (ThermoFisher Scientific). RNA with intact 28S and 18S ribosomal RNA bands at a ratio of approximately 2:1 with A_260/280_ of ∼2 and A_260/230_ of 1.8 – 2.2 was used for reverse transcription (RT). When tissues were used as a source, the integrity of DNAse I-treated RNA was determined using the Agilent RNA 6000 Pico Kit and an Agilent 2100 Bioanalyzer (Agilent Technologies). RNA with RIN values > 4 was used for RT.

RT was performed with 400 ng RNA per 20 µL reaction using the qPCR cDNA Synthesis Kit (PCR Biosystems). cDNA was treated with RNase H (New England Biolabs) for 20 min at 37°C and diluted 1:5 prior to qPCR.

Primers for qPCR were synthesised by Merck and used at 200 nM final (sequences in Supplemental Table 1). Primer efficiencies were assessed by standard curve and specificity was confirmed by melt curve analysis and agarose gel electrophoresis. qPCR was performed using 2x qPCR BIO SyGreen Blue Mix Lo-ROX (PCR Biosystems) in triplicate 10 µL reactions in 384-well plates (Applied Biosystems). Amplification was performed on a QuantStudio™5 Real-Time PCR System (ThermoFisher Scientific) using the following cycling parameters: initial denaturation and polymerase activation at 95°C for 2 min; 40 cycles of 95°C for 5 s and 60°C for 30 s, and a subsequent melt step: 95°C for 15 s, 60°C for 60 s, 95°C for 15 s.

Data was analysed using the QuantStudio™ Design & Analysis Software. Mean C_q_ values of technical triplicates were normalised to the reference genes *RPS13* or *IPO8* and experimental controls using the δδC_q_ method^75^. Data is presented as average log_2_ fold-change ± SD except for *TFAM* mRNA, which is presented as fold change, and human tissue data, which is presented as 2^-δCq^.

### RNAseq of IFN-β treated human iCell microglia

iCell human microglia (FujiFilm CDI) were derived from peripheral blood mononuclear cells from a donor with no known disease phenotypes. 3 independent vials of iCell microglia were seeded at 5 x 10^4^ cells per well across 3 independent 96-well plates and treated the following day with vehicle or 5 ng/mL human IFN-β (R&D Systems). Cells were treated for 6 or 24 h prior to collection and stored at −80°C prior to RNA extraction.

For cell lysis, RNAdvance Cell v2 (Beckman Coulter) was added to iCell Microglia in 96-well plates on dry ice. Plates were warmed to room temperature, treated with Proteinase K and cells were lysed by gentle pipetting followed by incubation at room temperature for 30 min. RNA extraction was performed according to manufacturer’s protocol on a Biomek i7 Hybrid robotic workstation (Beckman Coulter). mRNA enrichment and sequencing library preparation was performed using KAPA mRNA HyperPrep Kit (Roche) according to the manufacturer’s protocol on a Tecan Fluent® liquid handler. Library quality was assessed on a fragment analyser instrument using SS NGS fragment kit (1-6,000 bp; Agilent). Sequencing was performed on NovaSeq 6000 platform using a 200 cycles SP Reagent Kit v1.5 (both Illumina) at a loading concentration of 1.85 nM. Sequencing generated 194.6 Gb of data with Q30 of over 91.6%. The data will be deposited in the NCBI Sequence Read Archive (SRA) and referenced upon acceptance of the manuscript (currently available on request).

### Transcriptomics data analysis

FastQC files were generated and trimmed with TrimGalore^76^ and quality review was performed by FastQC^77^ before and after quality trim. Reads were aligned with STAR aligner (version 2.6.1d, using the default settings) to the HG38 human genome and ENSEMBL94 annotation GTF file. After alignment and summarization with featureCounts of the Subread package (feature Counts release 1.6.3 and picard version 2.18.21), samples were normalized using voom (limma package) and differential gene expression analysis was carried out with limma implemented in R. FDR adjusted p-value < 0.05 was used as the cut-off (Benjamini-Hochberg method). Gene annotations were obtained by biomaRt.

### Pathway and network analyses

For protein interaction network analysis and visualization Cytoscape was used (version 3.8.0) and the StringApp plugin using a confidence score cutoff of 0.7 (high confidence)^78,79^. Gene Ontology (GO) and KEGG database enrichment was performed using R and the clusterProfiler package with an adjusted p-value cutoff of 0.05^80^.

### Measurements of cytokine/chemokine secretion

Following 72 h pre-treatment with 10 nM AZD1390 or DMSO as a control, WT and *STING1* KO HMC3 cells were seeded at a density of 21,000 cells/cm^2^ in 6-well plates. After 48 h, cells were washed in PBS and placed in DMEM supplemented with 0.5% (v/v) HI-FBS. Cells were stimulated with 1 µM diABZi for 24 h prior to collection. Conditioned medium was centrifuged at 400 *g* for 10 min at 4°C to pellet cells and debris, then snap frozen on dry ice and stored at −80°C until use. After collection of medium, remaining cells were lysed in RIPA buffer and total protein content was measured by Bradford assay. The overall protein content was comparable between treatments.

Secretion of IFN-α2a, IFN-β, and CXCL10 was measured using a Mesoscale Discovery (MSD) U-PLEX assay following the manufacturer’s protocol. Plates were read on a MESO SECTOR S 600 (MSD) and data was analysed using MSD Discovery Workbench (version 4).

### Immunofluorescence

Cells were seeded on glass coverslips and fixed in 4% (w/v) paraformaldehyde for 15 min at room temperature. Coverslips were then permeabilised with 0.2% (v/v) Triton X-100 for 10 min and blocked in 3% (w/v) BSA, 0.05% (v/v) Tween®20, 0.3 M glycine in PBS for 1 h. For detection of IRF3, coverslips were fixed and permeabilised in ice-cold methanol for 5 min before blocking. Antibodies were diluted in blocking buffer (Supplemental Table 1) and cells were incubated with primary antibodies at 4°C overnight in a humidified environment. Coverslips were then washed in PBS with 0.1% (v/v) Tween®20 (PBST) and incubated with secondary antibodies for 45 min at room temperature. Coverslips were washed in PBST before mounting onto glass slides using Fluoroshield™ mounting medium (Merck) supplemented with 1.5 µg/mL DAPI.

Detection of cytosolic dsDNA was performed as previously described with amendments^81^. Coverslips were PFA-fixed and permeabilised with 0.2% (v/v) Triton X-100 for 5 min at 4°C. To remove cytosolic RNA, coverslips were treated with 2 mg/ml RNase A (QIAGEN) for 30 min at 37°C, then washed in PBS. Antibody specificity was validated by treating with 200 U/mL DNase I (ThermoFisher) for 30 min at 37°C. Coverslips were blocked in 3% (w/v) BSA, 0.3 M glycine in PBS. Coverslips were then processed as described above, however Triton X-100 was omitted from subsequent buffers to limit permeabilization of the nuclear membrane.

### Microscopy and Image Analysis

Fluorescence microscopy was performed using a Zeiss Axio Imager M2 or Zeiss Axio Imager Z2 epifluorescent microscope using a 20x objective. Laser scanning confocal microscopy was performed using a Nikon Eclipse Ti microscope with 40x or 60x oil-immersion objectives.

Images were processed and analysed using FIJI (ImageJ; National Institutes of Health) or CellProfiler (Broad Institute). To analyse nuclear translocation of IRF3, the nuclei and cytoplasm were segmented in CellProfiler using DAPI and vinculin staining, respectively. Mean fluorescence intensity was determined in each compartment and expressed as a ratio of nuclear:cytoplasmic (N:C) intensity. Cells with nuclear translocation were determined as the percentage of cells with an N:C ratio > 1.5.

### Quantification of micronuclei (MN)

Cells were grown on glass coverslips, fixed and DAPI-stained as above. Blinded MN scoring was performed using the ImageJ Cell Counter plugin in accordance with published criteria^82^. Mitotic and apoptotic cells were manually excluded from analysis. For detection of cGAS-positive MN, an intensity threshold was set, and cGAS-positive MN were identified by particle analysis in ImageJ. The number of cGAS-positive MN was normalised to the number of cells scored (> 100 cells per independent experiment).

### Quantification of cytoplasmic dsDNA

Cytoplasmic dsDNA intensity was analysed on maximum intensity projections of confocal Z-stacks using CellProfiler. Nuclei were segmented based on DAPI staining. The resulting nuclei and TOM20 staining were then used to define the whole-cell boundary and the nuclear region was subtracted from the whole-cell region to define the cytoplasm. Mean fluorescence intensity of the dsDNA signal was then measured in the cytoplasmic compartment. To determine the percentage of cytoplasmic dsDNA-positive cells, dsDNA speckles were enhanced to enable identification as primary objects. The resulting objects were filtered based on mean fluorescence intensity and to improve specificity, cells with at least 10 dsDNA foci were classified as positive for cytoplasmic dsDNA. Identification of the nucleus and consequently the whole cell boundary was reliant on DAPI intensity. As DNase I treatment abolished the DAPI signal, quantification of cytoplasmic intensity could not be performed for DNase I-treated conditions. Five fields of view were analysed per condition.

### Wound healing assay

WT and *STING1* KO HMC3 cells were pre-treated with 10 nM AZD1390 or DMSO as a control for 72 h, then seeded at a density of 79,000 cells/cm^2^ into 2-well culture inserts (Ibidi). After 24 h, inserts were removed leaving a 500 µm gap, cells were washed twice in PBS then placed in DMEM supplemented with 0.5% HI-FBS to limit proliferation. Wounds were imaged every 4 h over a period of 24 h on a Lumascope LS720 (Etaluma Inc) using a 4x phase-contrast objective. Wound closure was analysed in ImageJ and wound area is presented as a percentage of the 0-h time point.

### xCELLigence chemotaxis assay

For preparation of conditioned medium, WT and *STING1* KO HMC3 cells were pre-treated with 10 nM AZD1390 or DMSO as a control for 72 h, then seeded at a density of 21,000 cells/cm^2^ in 6-well plates. After 48 h, cells were washed in PBS and placed in DMEM supplemented with 0.5% (v/v) HI-FBS. Conditioned medium was centrifuged at 400 *g* for 10 min to remove whole cells and debris.

Chemotaxis assays were performed using Real-Time Cell Analysis (RTCA) CIM-16 plates (Agilent). The lower chamber contained conditioned medium as the chemoattractant or DMEM supplemented with 0.5% (v/v) HI-FBS as a negative control. The upper chamber was filled with 0.5% (v/v) HI-FBS and plates were left to equilibrate at 37°C for 1 h. Background impedance (Cell Index) was measured followed by seeding 25,000 cells per well in the upper chamber in technical triplicates. Cells were left to settle at RT for 30 min, then impedance was measured every 15 min for 6 h using an xCELLigence RTCA DP instrument (Agilent). Due to the nature of the assay, absolute cell index values differed between independent experiments precluding presentation of the summarised data.

### Statistical Analysis

Statistical analysis was performed using GraphPad Prism version 9.4.1. Quantitative data is presented as mean ± standard deviation (SD) from at least 3 independent experiments unless stated otherwise. Normality was assessed using Q-Q plots and the Shapiro-Wilk normality test. Student’s or Welch’s t-tests were used to compare means of two groups with equal or unequal variance, respectively. For comparison of 3 or more groups, one- or two-way ANOVAs were used. P-values ≤ 0.05 were deemed to be statistically significant.

## Acknowledgements

We thank all members of the S.V.K. laboratory for the discussion and especially Wen Cheng and Chandika Soondram for occasional help at the bench. We are grateful to Joana Cerveira at the Flow Cytometry Facility (Department of Pathology, University of Cambridge, UK) for discussions. We thank Ramy Elgendy (AstraZeneca) for contributing to RNA sequencing work. We thank Lucia Pereira Giraldez and Robert Harvey (MRC Toxicology Unit, University of Cambridge, UK) for providing access to the xCELLigence system and advice. We also thank Tsveta Kamenova (laboratory of Eric Miska, Department of Biochemistry, University of Cambridge, UK) for technical advice on tissue quality control. Human tissue was obtained from the NIH Neurobiobank at the University of Maryland, Baltimore, MD (United States).

The article is based on E.J.T.’s PhD thesis work^83^.

## Author contributions

E.J.T. and S.V.K. conceived the study and wrote the manuscript with contributions from L.J. and K.T. E.J.T. performed most of the experiments with the following exceptions. L.J. performed experiments with DNA damaging agents, tissue samples and chemotaxis assays. S.V.K. helped with cell line generation and cell synchronisation experiments. P.T. did the work with iCell microglia and advised on the use of the MSD secretion assay platform. M.D. performed RNA sequencing. K.T. performed bioinformatic analyses of RNA sequencing data. M.P. provided access to the MSD platform. All authors contributed to the data discussion. S.V.K. supervised the work.

## REFERENCES

1. Lin, X., Kapoor, A., Gu, Y., Chow, M.J., Peng, J., Zhao, K., and Tang, D. (2020). Contributions of DNA Damage to Alzheimer’s disease. Int. J. Mol. Sci. 21, 1666. 10.3390/ijms21051666.

2. Sun, Y., Curle, A.J., Haider, A.M., and Balmus, G. (2020). The role of DNA damage response in amyotrophic lateral sclerosis. Essays Biochem. 64, 847–861. 10.1042/EBC20200002.

3. López-Otín, C., Blasco, M.A., Partridge, L., Serrano, M., and Kroemer, G. (2023). Hallmarks of aging: An expanding universe. Cell 186, 243–278. 10.1016/j.cell.2022.11.001.

4. Welch, G., and Tsai, L.-H. (2022). Mechanisms of DNA damage-mediated neurotoxicity in neurodegenerative disease. EMBO Rep. 23, e54217. 10.15252/embr.202154217.

5. Aditi, and McKinnon, P.J. (2022). Genome integrity and inflammation in the nervous system. DNA Repair 119, 103406. 10.1016/j.dnarep.2022.103406.

6. Härtlova, A., Erttmann, S.F., Raffi, F.A.M., Schmalz, A.M., Resch, U., Anugula, S., Lienenklaus, S., Nilsson, L.M., Kröger, A., Nilsson, J.A., et al. (2015). DNA damage primes the type I interferon system via the cytosolic DNA sensor STING to promote anti-microbial innate immunity. Immunity 42, 332–343. 10.1016/j.immuni.2015.01.012.

7. Decout, A., Katz, J.D., Venkatraman, S., and Ablasser, A. (2021). The cGAS–STING pathway as a therapeutic target in inflammatory diseases. Nat. Rev. Immunol. 21, 548–569. 10.1038/s41577-021-00524-z.

8. Hopfner, K.-P., and Hornung, V. (2020). Molecular mechanisms and cellular functions of cGAS–STING signalling. Nat. Rev. Mol. Cell Biol. 21, 501–521. 10.1038/s41580-020-0244-x.

9. Harding, S.M., Benci, J.L., Irianto, J., Discher, D.E., Minn, A.J., and Greenberg, R.A. (2017). Mitotic progression following DNA damage enables pattern recognition within micronuclei. Nature 548, 466–470.

10. Mackenzie, K.J., Carroll, P., Martin, C.-A., Murina, O., Fluteau, A., Simpson, D.J., Olova, N., Sutcliffe, H., Rainger, J.K., Leitch, A., et al. (2017). cGAS surveillance of micronuclei links genome instability to innate immunity. Nature 548, 461–465.

11. MacDonald, K.M., Nicholson-Puthenveedu, S., Tageldein, M.M., Khasnis, S., Arrowsmith, C.H., and Harding, S.M. (2023). Antecedent chromatin organization determines cGAS recruitment to ruptured micronuclei. Nat. Commun. 14, 556. 10.1038/s41467-023-36195-8.

12. Song, X., Aw, J.T.M., Ma, F., Cheung, M.F., Leung, D., and Herrup, K. (2021). DNA repair inhibition leads to active export of repetitive sequences to the cytoplasm triggering an inflammatory response. J. Neurosci. 41, 9286–9307. 10.1523/JNEUROSCI.0845-21.2021.

13. West, A.P., Khoury-Hanold, W., Staron, M., Tal, M.C., Pineda, C.M., Lang, S.M., Bestwick, M., Duguay, B.A., Raimundo, N., MacDuff, D.A., et al. (2015). Mitochondrial DNA stress primes the antiviral innate immune response. Nature 520, 553–557. 10.1038/nature14156.

14. Crossley, M.P., Song, C., Bocek, M.J., Choi, J.-H., Kousorous, J., Sathirachinda, A., Lin, C., Brickner, J.R., Bai, G., Lans, H., et al. (2023). R-loop-derived cytoplasmic RNA–DNA hybrids activate an immune response. Nature 613, 187–194. 10.1038/s41586-022-05545-9.

15. Chen, Y.-A., Shen, Y.-L., Hsia, H.-Y., Tiang, Y.-P., Sung, T.-L., and Chen, L.-Y. (2017). Extrachromosomal telomere repeat DNA is linked to ALT development via cGAS-STING DNA sensing pathway. Nat. Struct. Mol. Biol. 24, 1124–1131. 10.1038/nsmb.3498.

16. Reinert, L.S., Lopušná, K., Winther, H., Sun, C., Thomsen, M.K., Nandakumar, R., Mogensen, T.H., Meyer, M., Vægter, C., Nyengaard, J.R., et al. (2016). Sensing of HSV-1 by the cGAS–STING pathway in microglia orchestrates antiviral defence in the CNS. Nat. Commun. 7, 13348.

17. Paolicelli, R.C., Bolasco, G., Pagani, F., Maggi, L., Scianni, M., Panzanelli, P., Giustetto, M., Ferreira, T.A., Guiducci, E., Dumas, L., et al. (2011). Synaptic pruning by microglia is necessary for normal brain development. Science 333, 1456–1458. 10.1126/science.1202529.

18. Schafer, D.P., Lehrman, E.K., Kautzman, A.G., Koyama, R., Mardinly, A.R., Yamasaki, R., Ransohoff, R.M., Greenberg, M.E., Barres, B.A., and Stevens, B. (2012). Microglia sculpt postnatal neural circuits in an activity and complement-dependent manner. Neuron 74, 691–705. 10.1016/j.neuron.2012.03.026.

19. Bellver-Landete, V., Bretheau, F., Mailhot, B., Vallières, N., Lessard, M., Janelle, M.-E., Vernoux, N., Tremblay, M.-È., Fuehrmann, T., Shoichet, M.S., et al. (2019). Microglia are an essential component of the neuroprotective scar that forms after spinal cord injury. Nat. Commun. 10, 518. 10.1038/s41467-019-08446-0.

20. Sierra, A., Encinas, J.M., Deudero, J.J.P., Chancey, J.H., Enikolopov, G., Overstreet-Wadiche, L.S., Tsirka, S.E., and Maletic-Savatic, M. (2010). Microglia shape adult hippocampal neurogenesis through apoptosis-coupled phagocytosis. Cell Stem Cell 7, 483–495. 10.1016/j.stem.2010.08.014.

21. Muzio, L., Viotti, A., and Martino, G. (2021). Microglia in neuroinflammation and neurodegeneration: from understanding to therapy. Front. Neurosci. 15, 742065.

22. Hickman, S., Izzy, S., Sen, P., Morsett, L., and Khoury, J.E. (2018). Microglia in neurodegeneration. Nat. Neurosci. 21, 1359–1369. 10.1038/s41593-018-0242-x.

23. Shiloh, Y., and Lederman, H.M. (2017). Ataxia-telangiectasia (A-T): an emerging dimension of premature ageing. Ageing Res. Rev. 33, 76–88. 10.1016/j.arr.2016.05.002.

24. Bourseguin, J., Cheng, W., Talbot, E., Hardy, L., Lai, J., Jeffries, A.M., Lodato, M.A., Lee, E.A., and Khoronenkova, S.V. (2022). Persistent DNA damage associated with ATM kinase deficiency promotes microglial dysfunction. Nucleic Acids Res. 50, 2700–2718. 10.1093/nar/gkac104.

25. Quek, H., Luff, J., Cheung, K.G., Kozlov, S., Gatei, M., Lee, C.S., Bellingham, M.C., Noakes, P.G., Lim, Y.C., Barnett, N.L., et al. (2017). A rat model of ataxia-telangiectasia: evidence for a neurodegenerative phenotype. Hum. Mol. Genet. 26, 109–123. 10.1093/hmg/ddw371.

26. Song, X., Ma, F., and Herrup, K. (2019). Accumulation of cytoplasmic DNA due to ATM deficiency activates the microglial viral response system with neurotoxic consequences. J. Neurosci. 39, 6378–6394. 10.1523/JNEUROSCI.0774-19.2019.

27. Levi, H., Bar, E., Cohen-Adiv, S., Sweitat, S., Kanner, S., Galron, R., Mitiagin, Y., and Barzilai, A. (2022). Dysfunction of cerebellar microglia in ataxia-telangiectasia. Glia 70, 536–557. 10.1002/glia.24122.

28. Quek, H., Luff, J., Cheung, K., Kozlov, S., Gatei, M., Lee, C.S., Bellingham, M.C., Noakes, P.G., Lim, Y.C., Barnett, N.L., et al. (2016). Rats with a missense mutation in Atm display neuroinflammation and neurodegeneration subsequent to accumulation of cytosolic DNA following unrepaired DNA damage. J. Leukoc. Biol. 101, 927–947. 10.1189/jlb.4vma0716-316r.

29. Grabert, K., Michoel, T., Karavolos, M.H., Clohisey, S., Baillie, J.K., Stevens, M.P., Freeman, T.C., Summers, K.M., and McColl, B.W. (2016). Microglial brain region−dependent diversity and selective regional sensitivities to aging. Nat. Neurosci. 19, 504–516. 10.1038/nn.4222.

30. Tay, T.L., Mai, D., Dautzenberg, J., Fernández-Klett, F., Lin, G., Sagar, Datta, M., Drougard, A., Stempfl, T., Ardura-Fabregat, A., et al. (2017). A new fate mapping system reveals context-dependent random or clonal expansion of microglia. Nat. Neurosci. 20, 793–803. 10.1038/nn.4547.

31. Lai, J., Kim, J., Jeffries, A.M., Tolles, A., Chittenden, T.W., Buckley, P.G., Yu, T.W., Lodato, M.A., and Lee, E.A. (2021). Single-nucleus transcriptomic analyses reveal microglial activation underlying cerebellar degeneration in Ataxia Telangiectasia.

32. Aguado, J., Chaggar, H.K., Gómez-Inclán, C., Shaker, M.R., Leeson, H.C., Mackay-Sim, A., and Wolvetang, E.J. (2021). Inhibition of the cGAS-STING pathway ameliorates the premature senescence hallmarks of Ataxia-Telangiectasia brain organoids. Aging Cell 20, e13468. 10.1111/acel.13468.

33. Hu, M., Zhou, M., Bao, X., Pan, D., Jiao, M., Liu, X., Li, F., and Li, C.-Y. (2021). ATM inhibition enhances cancer immunotherapy by promoting mtDNA leakage and cGAS/STING activation. J. Clin. Invest. 131, e139333. 10.1172/JCI139333.

34. Zhang, Q., Green, M.D., Lang, X., Lazarus, J., Parsels, J.D., Wei, S., Parsels, L.A., Shi, J., Ramnath, N., Wahl, D.R., et al. (2019). Inhibition of ATM increases interferon signaling and sensitizes pancreatic cancer to immune checkpoint blockade therapy. Cancer Res. 79, 3940–3951. 10.1158/0008-5472.CAN-19-0761.

35. Yang, B., Dan, X., Hou, Y., Lee, J.-H., Wechter, N., Krishnamurthy, S., Kimura, R., Babbar, M., Demarest, T., McDevitt, R., et al. (2021). NAD+ supplementation prevents STING-induced senescence in ataxia telangiectasia by improving mitophagy. Aging Cell 20, e13329. 10.1111/acel.13329.

36. Concannon, P., and Gatti, R.A. (1997). Diversity of ATM gene mutations detected in patients with ataxia-telangiectasia. Hum. Mutat. 10, 100–107. 10.1002/(SICI)1098-1004(1997)10:2<100::AID-HUMU2>3.0.CO;2-O.

37. Sakasai, R., Teraoka, H., Takagi, M., and Tibbetts, R.S. (2010). Transcription-dependent activation of ataxia telangiectasia mutated prevents DNA-dependent protein kinase-mediated cell death in response to topoisomerase I poison. J. Biol. Chem. 285, 15201– 15208. 10.1074/jbc.M110.101808.

38. Ramanjulu, J.M., Pesiridis, G.S., Yang, J., Concha, N., Singhaus, R., Zhang, S.-Y., Tran, J.-L., Moore, P., Lehmann, S., Eberl, H.C., et al. (2018). Design of amidobenzimidazole STING receptor agonists with systemic activity. Nature 564, 439–443. 10.1038/s41586-018-0705-y.

39. Durant, S.T., Zheng, L., Wang, Y., Chen, K., Zhang, L., Zhang, T., Yang, Z., Riches, L., Trinidad, A.G., Fok, J.H.L., et al. (2018). The brain-penetrant clinical ATM inhibitor AZD1390 radiosensitizes and improves survival of preclinical brain tumor models. Sci. Adv. 4, eaat1719. 10.1126/sciadv.aat1719.

40. Trifonova, R.T., and Barteneva, N.S. (2018). Quantitation of IRF3 nuclear translocation in heterogeneous cellular populations from cervical tissue using imaging flow cytometry. In Cellular Heterogeneity: Methods and Protocols Methods in Molecular Biology., N. S. Barteneva and I. A. Vorobjev, eds. (Springer), pp. 125–153.

41. Gómez-Nicola, D., Fransen, N.L., Suzzi, S., and Perry, V.H. (2013). Regulation of microglial proliferation during chronic neurodegeneration. J. Neurosci. 33, 2481–2493. 10.1523/JNEUROSCI.4440-12.2013.

42. Huang, Y., Xu, Z., Xiong, S., Sun, F., Qin, G., Hu, G., Wang, J., Zhao, L., Liang, Y.-X., Wu, T., et al. (2018). Repopulated microglia are solely derived from the proliferation of residual microglia after acute depletion. Nat. Neurosci. 21, 530–540. 10.1038/s41593-018-0090-8.

43. Yamamoto, K., Wang, J., Sprinzen, L., Xu, J., Haddock, C.J., Li, C., Lee, B.J., Loredan, D.G., Jiang, W., Vindigni, A., et al. (2016). Kinase-dead ATM protein is highly oncogenic and can be preferentially targeted by Topo-isomerase I inhibitors. eLife 5, e14709. 10.7554/eLife.14709.

44. Yamamoto, K., Wang, Y., Jiang, W., Liu, X., Dubois, R.L., Lin, C.-S., Ludwig, T., Bakkenist, C.J., and Zha, S. (2012). Kinase-dead ATM protein causes genomic instability and early embryonic lethality in mice. J. Cell Biol. 198, 305–313. 10.1083/jcb.201204098.

45. Kranzusch, P.J., Lee, A.S.-Y., Berger, J.M., and Doudna, J.A. (2013). Structure of Human cGAS Reveals a Conserved Family of Second-Messenger Enzymes in Innate Immunity. Cell Rep. 3, 1362–1368. 10.1016/j.celrep.2013.05.008.

46. Valentin-Vega, Y.A., MacLean, K.H., Tait-Mulder, J., Milasta, S., Steeves, M., Dorsey, F.C., Cleveland, J.L., Green, D.R., and Kastan, M.B. (2012). Mitochondrial dysfunction in ataxia-telangiectasia. Blood 119, 1490–1500. 10.1182/blood-2011-08-373639.

47. Pillai, S., Nguyen, J., Johnson, J., Haura, E., Coppola, D., and Chellappan, S. (2015). Tank binding kinase 1 is a centrosome-associated kinase necessary for microtubule dynamics and mitosis. Nat. Commun. 6, 10072. 10.1038/ncomms10072.

48. Sarraf, S.A., Sideris, D.P., Giagtzoglou, N., Ni, L., Kankel, M.W., Sen, A., Bochicchio, L.E., Huang, C.-H., Nussenzweig, S.C., Worley, S.H., et al. (2019). PINK1/Parkin influences cell cycle by sequestering TBK1 at damaged mitochondria, inhibiting mitosis. Cell Rep. 29, 225–235.e5. 10.1016/j.celrep.2019.08.085.

49. Dunphy, G., Flannery, S.M., Almine, J.F., Connolly, D.J., Paulus, C., Jønsson, K.L., Jakobsen, M.R., Nevels, M.M., Bowie, A.G., and Unterholzner, L. (2018). Non-canonical activation of the DNA sensing adaptor STING by ATM and IFI16 mediates NF-κB signaling after nuclear DNA damage. Mol. Cell 71, 745–760.e5. 10.1016/j.molcel.2018.07.034.

50. Chu, T.-T., Tu, X., Yang, K., Wu, J., Repa, J.J., and Yan, N. (2021). Tonic prime-boost of STING signalling mediates Niemann–Pick disease type C. Nature 596, 570–575. 10.1038/s41586-021-03762-2.

51. de Oliveira Mann, C.C., Orzalli, M.H., King, D.S., Kagan, J.C., Lee, A.S.Y., andKranzusch, P.J. (2019). Modular architecture of the STING C-terminal tail allows interferon and NF-κB signaling adaptation. Cell Rep. 27, 1165–1175.e5. 10.1016/j.celrep.2019.03.098.

52. Hughes, C.E., and Nibbs, R.J.B. (2018). A guide to chemokines and their receptors. Febs J. 285, 2944–2971. 10.1111/febs.14466.

53. Holm, C.K., Jensen, S.B., Jakobsen, M.R., Cheshenko, N., Horan, K.A., Moeller, H.B., Gonzalez-Dosal, R., Rasmussen, S.B., Christensen, M.H., Yarovinsky, T.O., et al. (2012). Virus-cell fusion as a trigger of innate immunity dependent on the adaptor STING. Nat. Immunol. 13, 737–743. 10.1038/ni.2350.

54. Marques, J.T., and Williams, B.R.G. (2005). Activation of the mammalian immune system by siRNAs. Nat. Biotechnol. 23, 1399–1405. 10.1038/nbt1161.

55. Nimmerjahn, A., Kirchhoff, F., and Helmchen, F. (2005). Resting microglial cells are highly dynamic surveillants of brain parenchyma in vivo. Science 308, 1314–1318. 10.1126/science.1110647.

56. Limame, R., Wouters, A., Pauwels, B., Fransen, E., Peeters, M., Lardon, F., Wever, O.D., and Pauwels, P. (2012). Comparative Analysis of Dynamic Cell Viability, Migration and Invasion Assessments by Novel Real-Time Technology and Classic Endpoint Assays. PLOS ONE 7, e46536. 10.1371/journal.pone.0046536.

57. Balmus, G., Pilger, D., Coates, J., Demir, M., Sczaniecka-Clift, M., Barros, A.C., Woods, M., Fu, B., Yang, F., Chen, E., et al. (2019). ATM orchestrates the DNA-damage response to counter toxic non-homologous end-joining at broken replication forks. Nat. Commun. 10, 87. 10.1038/s41467-018-07729-2.

58. Wilson, D.M., Cookson, M.R., Van Den Bosch, L., Zetterberg, H., Holtzman, D.M., and Dewachter, I. (2023). Hallmarks of neurodegenerative diseases. Cell 186, 693–714. 10.1016/j.cell.2022.12.032.

59. Aw, E., Zhang, Y., and Carroll, M. (2020). Microglial responses to peripheral type 1 interferon. J. Neuroinflammation 17, 340. 10.1186/s12974-020-02003-z.

60. Li, W., Viengkhou, B., Denyer, G., West, P.K., Campbell, I.L., and Hofer, M.J. (2018). Microglia have a more extensive and divergent response to interferon-a compared with astrocytes. Glia 66, 2058–2078. 10.1002/glia.23460.

61. Rappert, A. (2004). CXCR3-Dependent Microglial Recruitment Is Essential for Dendrite Loss after Brain Lesion. J. Neurosci. 24, 8500–8509. 10.1523/JNEUROSCI.2451-04.2004.

62. Festa, B.P., Siddiqi, F.H., Jimenez-Sanchez, M., Won, H., Rob, M., Djajadikerta, A., Stamatakou, E., and Rubinsztein, D.C. (2023). Microglial-to-neuronal CCR5 signaling regulates autophagy in neurodegeneration. Neuron 0. 10.1016/j.neuron.2023.04.006.

63. Haruwaka, K., Ikegami, A., Tachibana, Y., Ohno, N., Konishi, H., Hashimoto, A., Matsumoto, M., Kato, D., Ono, R., Kiyama, H., et al. (2019). Dual microglia effects on blood brain barrier permeability induced by systemic inflammation. Nat. Commun. 10, 5816. 10.1038/s41467-019-13812-z.

64. Chen, X., Firulyova, M., Manis, M., Herz, J., Smirnov, I., Aladyeva, E., Wang, C., Bao, X., Finn, M.B., Hu, H., et al. (2023). Microglia-mediated T cell infiltration drives neurodegeneration in tauopathy. Nature 615, 668–677. 10.1038/s41586-023-05788-0.

65. Anastasiou, M., Newton, G.A., Kaur, K., Carrillo-Salinas, F.J., Smolgovsky, S.A., Bayer, A.L., Ilyukha, V., Sharma, S., Poltorak, A., Luscinskas, F.W., et al. (2021). Endothelial STING controls T cell transmigration in an IFNI-dependent manner. JCI Insight 6, e149346. 10.1172/jci.insight.149346.

66. Nagata, M., Kosaka, A., Yajima, Y., Yasuda, S., Ohara, M., Ohara, K., Harabuchi, S., Hayashi, R., Funakoshi, H., Ueda, J., et al. (2021). A critical role of STING-triggered tumor-migrating neutrophils for anti-tumor effect of intratumoral cGAMP treatment. Cancer Immunol. Immunother. 70, 2301–2312. 10.1007/s00262-021-02864-0.

67. Tan, J., Ge, Y., Zhang, M., and Ding, M. (2023). Proteomics analysis uncovers plasminogen activator PLAU as a target of the STING pathway for suppression of cancer cell migration and invasion. J. Biol. Chem. 299. 10.1016/j.jbc.2022.102779.

68. Tolbert, C.E., Beck, M.V., Kilmer, C.E., and Srougi, M.C. (2019). Loss of ATM positively regulates Rac1 activity and cellular migration through oxidative stress. Biochem. Biophys. Res. Commun. 508, 1155–1161. 10.1016/j.bbrc.2018.12.033.

69. Stowell, R.D., Wong, E.L., Batchelor, H.N., Mendes, M.S., Lamantia, C.E., Whitelaw, B.S., and Majewska, A.K. (2018). Cerebellar microglia are dynamically unique and survey Purkinje neurons in vivo. Dev. Neurobiol. 78, 627–644. 10.1002/dneu.22572.

70. Gulen, M.F., Samson, N., Keller, A., Schwabenland, M., Liu, C., Glück, S., Thacker, V.V., Favre, L., Mangeat, B., Kroese, L.J., et al. (2023). cGAS–STING drives ageing-related inflammation and neurodegeneration. Nature, 1–7. 10.1038/s41586-023-06373-1.

71. Janabi, N., Peudenier, S., Héron, B., Ng, K.H., and Tardieu, M. (1995). Establishment of human microglial cell lines after transfection of primary cultures of embryonic microglial cells with the SV40 large T antigen. Neurosci. Lett. 195, 105–108. 10.1016/0304-3940(94)11792-H.

72. Garcia-Mesa, Y., Jay, T.R., Checkley, M.A., Luttge, B., Dobrowolski, C., Valadkhan, S., Landreth, G.E., Karn, J., and Alvarez-Carbonell, D. (2017). Immortalization of primary microglia: a new platform to study HIV regulation in the central nervous system. J. Neurovirol. 23, 47–66. 10.1007/s13365-016-0499-3.

73. Ran, F.A., Hsu, P.D., Wright, J., Agarwala, V., Scott, D.A., and Zhang, F. (2013). Genome engineering using the CRISPR-Cas9 system. Nat. Protoc. 8, 2281–2308. 10.1038/nprot.2013.143.

74. Botling, J., and Micke, P. (2011). Fresh frozen tissue: RNA extraction and quality control. In Methods in Biobanking Methods in Molecular Biology., J. Dillner, ed. (Humana Press), pp. 405–413.

75. Livak, K.J., and Schmittgen, T.D. (2001). Analysis of relative gene expression data using real-time quantitative PCR and the 2-ΔΔCT method. Methods 25, 402–408. 10.1006/meth.2001.1262.

76. Krueger, F., James, F., Ewels, P., Afyounian, E., and Schuster-Boeckler, B. (2021). TrimGalore: v0.6.7. 10.5281/zenodo.5127899.

77. Andrews, S. (2020). FastQC: v0.11.9. https://www.bioinformatics.babraham.ac.uk/projects/fastqc/.

78. Shannon, P., Markiel, A., Ozier, O., Baliga, N.S., Wang, J.T., Ramage, D., Amin, N., Schwikowski, B., and Ideker, T. (2003). Cytoscape: a software environment for integrated models of biomolecular interaction networks. Genome Res. 13, 2498–2504. 10.1101/gr.1239303.

79. Doncheva, N.T., Morris, J.H., Gorodkin, J., and Jensen, L.J. (2019). Cytoscape StringApp: Network Analysis and Visualization of Proteomics Data. J. Proteome Res. 18, 623–632. 10.1021/acs.jproteome.8b00702.

80. Wu, T., Hu, E., Xu, S., Chen, M., Guo, P., Dai, Z., Feng, T., Zhou, L., Tang, W., Zhan, L., et al. (2021). clusterProfiler 4.0: A universal enrichment tool for interpreting omics data. The Innovation 2, 100141. 10.1016/j.xinn.2021.100141.

81. Lam, A.R., Bert, N.L., Ho, S.S.W., Shen, Y.J., Tang, M.L.F., Xiong, G.M., Croxford, J.L., Koo, C.X., Ishii, K.J., Akira, S., et al. (2014). RAE1 ligands for the NKG2D receptor are regulated by STING-dependent DNA sensor pathways in lymphoma. Cancer Res. 74, 2193–2203. 10.1158/0008-5472.CAN-13-1703.

82. Fenech, M., Chang, W.P., Kirsch-Volders, M., Holland, N., Bonassi, S., and Zeiger, E. (2003). HUMN project: Detailed description of the scoring criteria for the cytokinesis-block micronucleus assay using isolated human lymphocyte cultures. Mutat. Res. - Genet. Toxicol. Environ. Mutagen. 534, 65–75. 10.1016/S1383-5718(02)00249-8.

83. Talbot, E. (2023). Mechanisms of cytosolic DNA sensing and neuroinflammation in ataxia-telangiectasia. [Apollo - University of Cambridge Repository]. 10.17863/CAM.97149.

