## Supplemental Information for "The cGAS-STING pathway regulates microglial chemotaxis in genome instability"

**Supplemental Figure 1:** ATM deficiency activates the cGAS-STING pathway in human microglia-like cells.

**Supplemental Figure 2:** Cytosolic DNA in ATM deficiency primarily derives from micronuclei.

**Supplemental Figure 3:** cGAS-STING drives type I IFN signalling upon loss of ATM activity in human microglia-like cells.

**Supplemental Figure 4:** STING drives expression and secretion of type I interferons following ATM inhibition in microglia-like cells.

**Supplemental Figure 5:** Persistent DNA damage drives STING-dependent chemokine induction in microglia-like cells.

**Supplemental Table 1:** Cell lines, reagents and software programs used in this work.

**Supplemental Table 2:** Human tissue sample information.

**Supplemental references.**

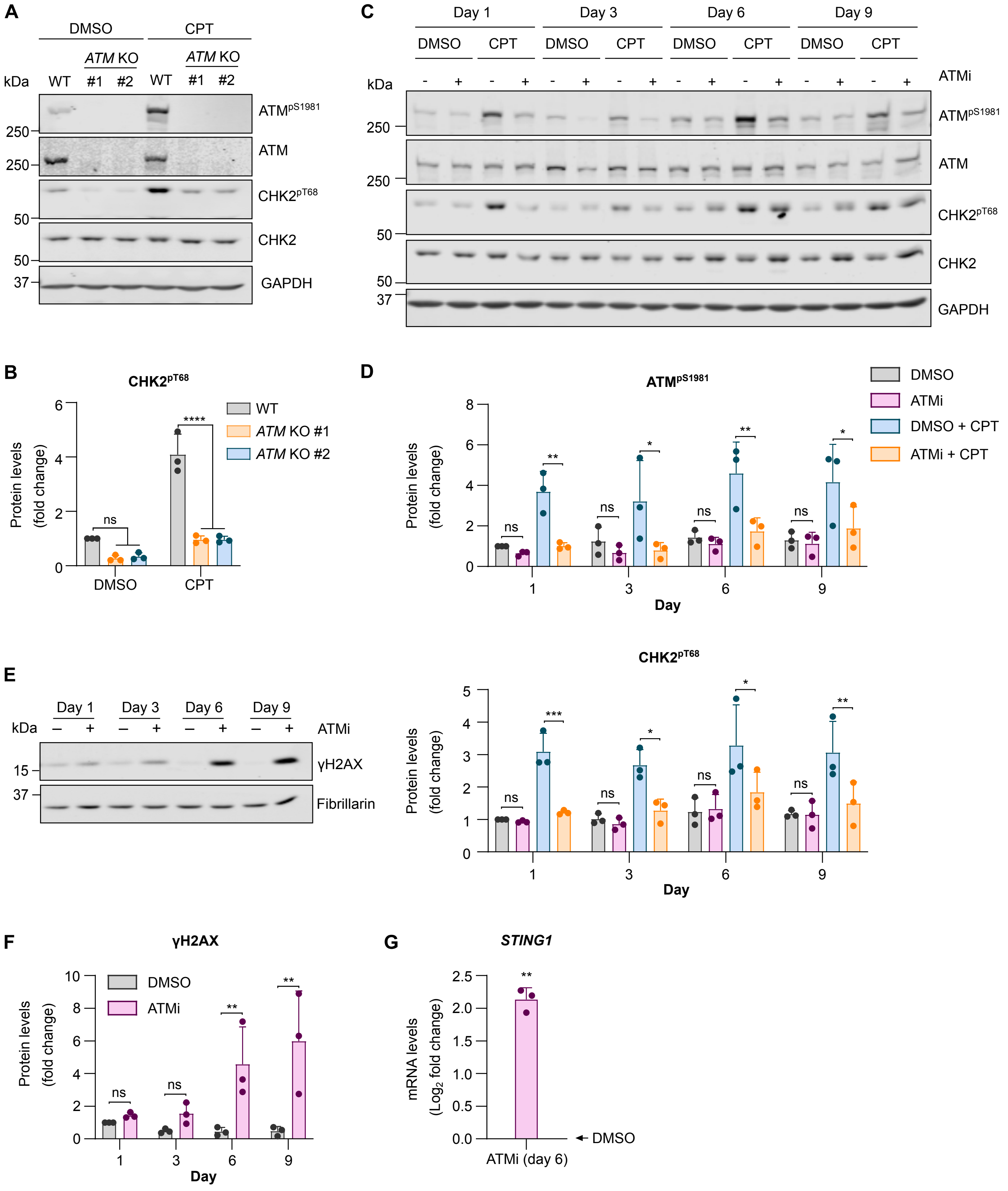


Talbot *et al.*, Supplemental Figure 1

**Supplemental Figure 1: ATM deficiency activates the cGAS-STING pathway in human microglia-like cells.**

**(A)** Representative immunoblot of *ATM* KO C20 microglial cells (clones #1 and #2) treated with 1 µM camptothecin (CPT), or DMSO as a control, for 1 h. Loading control: GAPDH.

**(B)** Relative quantification of $CHK2^{pT68}$ normalised to total CHK2 levels in C20 cells as in (A). Mean fold change ± SD (n=3). Two-way ANOVA with Tukey's post-hoc comparison test.

**(C)** Representative immunoblot of HMC3 cells treated as in Figure 1C. CPT or DMSO treatment as in (A). Loading control: GAPDH.

**(D)** Quantification of $ATM^{pS1981}$ and $CHK2^{pT68}$ normalised to respective total protein levels as in (C). Data are normalised to day 1 DMSO. Mean fold change ± SD (n=3). Two-way ANOVA with Tukey's post-hoc comparison test.

**(E)** Representative immunoblot of nuclear extracts of HMC3 cells treated as in (C). Loading control: Fibrillarin.

**(F)** Quantification of γH2AX normalised to Fibrillarin levels as in (E). Data are normalised to day 1 DMSO. Mean fold change ± SD (n=3). Two-way ANOVA with Sidak's multiple comparisons test.

**(G)** Relative *STING1* mRNA levels in HMC3 cells treated with ATMi for 6 days. $C_q$ values normalised to *RPS13* and DMSO-treated cells (indicated by arrow). Mean $log_2$ fold change ± SD (n=3). One-sample *t*-test comparing to a hypothetical mean of 0.

*p ≤ 0.05; **p ≤ 0.01; ***p ≤ 0.001; ****p ≤ 0.0001; ns = not significant.

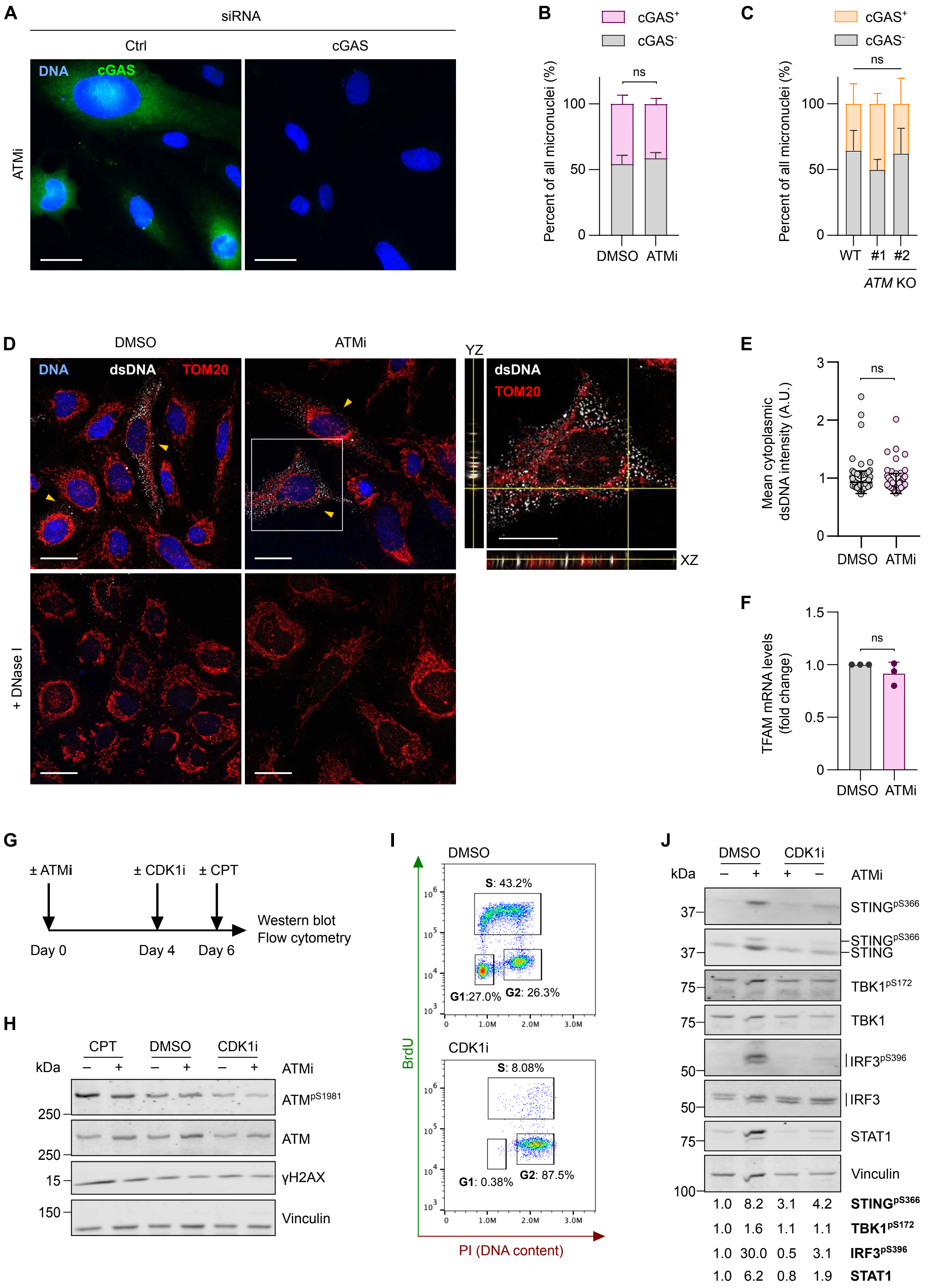


Talbot et al., Supplemental Figure 2

**Supplemental Figure 2. Cytosolic DNA in ATM deficiency primarily derives from micronuclei.**

**(A) I**mmunofluorescence microscopy images of HMC3 cells treated with 10 nM AZD1390 (ATMi) for 6 days in combination with Ctrl or cGAS siRNA. 20x objective, scale bar = 40 µm.
**(B)** Percentage of total MN positive or negative for cGAS as in Figure 2G. Mean ± SD (n=3). Two-way ANOVA with Tukey's post-hoc comparison test.
**(C)** Percentage of total MN positive or negative for cGAS as in Figure 2H. Mean ± SD (n=4). Two-way ANOVA with Tukey's post-hoc comparison test.
**(D)** Confocal microscopy images of HMC3 cells treated with 10 nM AZD1390 (ATMi), or DMSO as a control, for 6 days. Blue: DNA (DAPI), grey: dsDNA, red: TOM20 (mitochondria). Maximum intensity projections from Z-stacks, 40x oil immersion objective, scale bar = 15 µm. Yellow arrows indicate cells with cytosolic dsDNA. Inset: single Z-plane of highlighted ATMi-treated cell with YZ and XZ projections. Representative of 2 independent experiments.
**(E)** Quantification of cytoplasmic dsDNA intensity per cell as in (D). Mean intensity ± SD (n=1, at least 100 cells analysed per condition). Unpaired, two-tailed Student's t-test. A.U. = arbitrary units.
**(F)** Relative *TFAM* mRNA levels in HMC3 cells treated with ATMi or DMSO for 6 days. $C_q$ values normalised to *IPO8* and DMSO-treated cells. Mean $\log_2$ fold change ± SD (n=3). One-sample *t*-test, hypothetical mean of 1.
**(G)** Schematic of workflow. HMC3 cells were treated with 10 nM ATMi, or DMSO as a control, for 6 days in the presence of 9 µM RO-3306 (CDK1i) or DMSO for the final 48 h. To confirm ATM inhibition, cells were treated with 1 µM camptothecin (CPT) or DMSO for 1 h.
**(H)** Representative immunoblot of HMC3 cells treated as in (G). Loading control: Vinculin.
**(I)** Cells were treated as in (G), pulsed with 10 µM BrdU and analysed by BrdU-PI (propidium iodide) flow cytometry. Cell cycle distribution based on BrdU and PI intensity in DMSO (asynchronous) and CDK1i-treated cells (arrested in G2) in a representative experiment.
**(J)** Representative immunoblot of HMC3 cells treated with CDK1i, or DMSO as a control, as in (G). Loading control: Vinculin. Quantification of phosphorylated proteins and STAT1 is relative to respective total proteins and vinculin, respectively, and normalised to Day 6 DMSO control.
ns = not significant

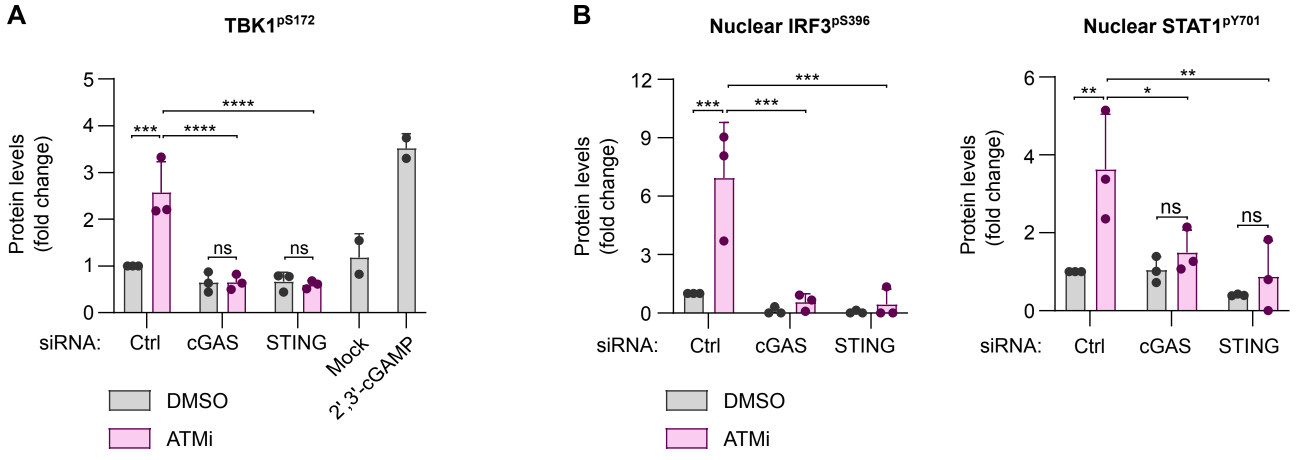


Talbot *et al.*, Supplemental Figure 3

**Supplemental Figure 3. cGAS-STING drives type I IFN signalling upon loss of ATM activity in human microglia-like cells.**

**(A)** Relative quantification of TBK1$^{pS172}$ as in Figure 3A normalised to total protein and DMSO- and control (Ctrl) siRNA-treated cells. Mean ± SD (n=3, except 2’,3’-cGAMP treatment where n=2). Two-way ANOVA with Tukey’s post-hoc comparison test.

**(B)** Relative quantification of IRF3$^{pS396}$ and STAT1$^{pY701}$ in nuclear extracts as in Figure 3B normalised to Fibrillarin and DMSO- and Ctrl siRNA-treated cells. Mean ± SD (n=3). Two-way ANOVA with Tukey’s post-hoc comparison test.

*p ≤ 0.05; **p ≤ 0.01; ***p ≤ 0.001; ****p ≤ 0.0001; ns = not significant.

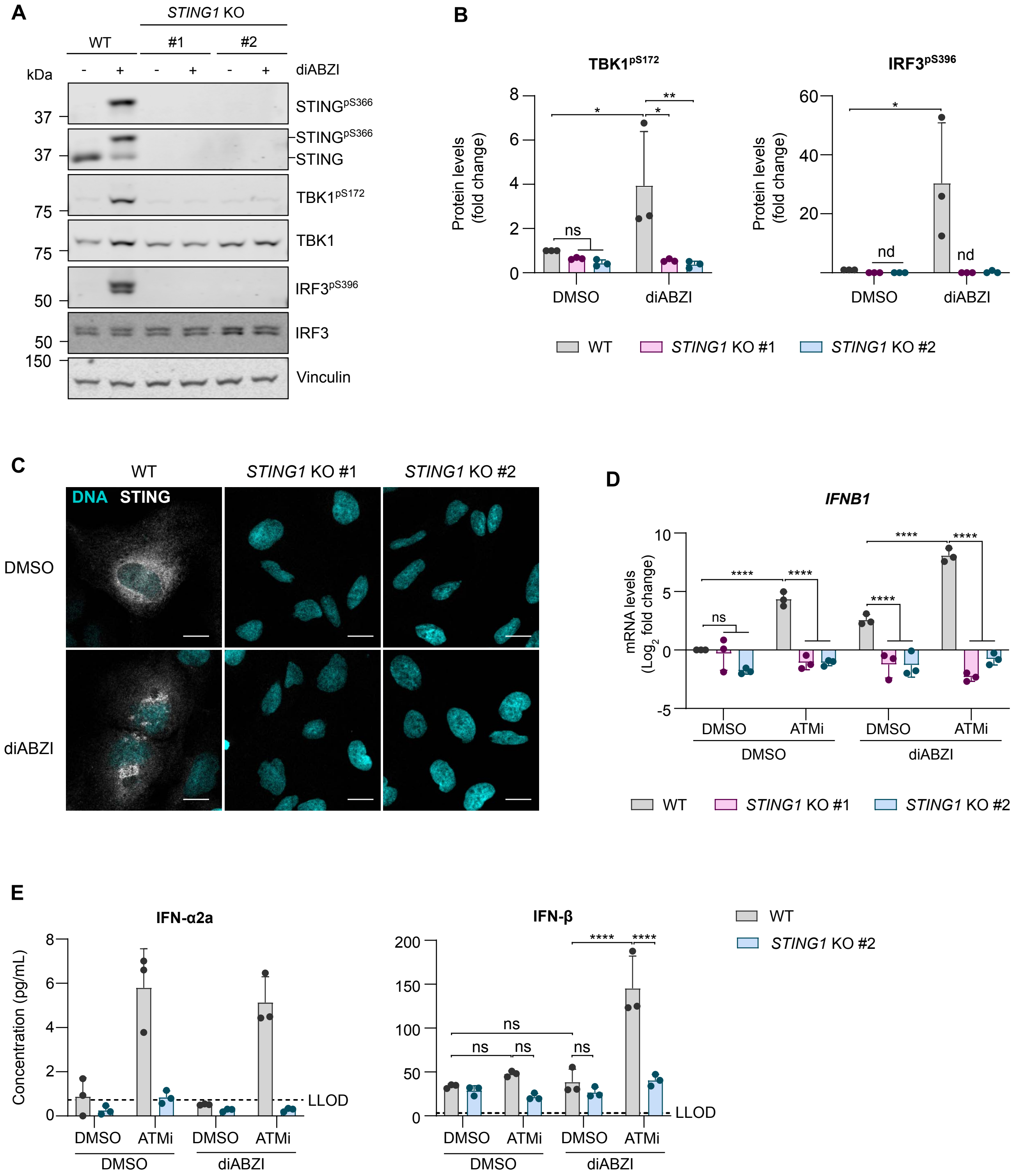


Talbot *et al.*, Supplemental Figure 4

**Supplemental Figure 4. STING drives expression and secretion of type I interferons following ATM inhibition in microglia-like cells.**

**(A)** Immunoblot of WT and *STING1* KO HMC3 cells (clones #1 and #2) treated with 1 µM diABZI, or DMSO as a control, for 2 h. Loading control: vinculin.

**(B)** Relative quantification of $TBK1^{pS172}$ and $IRF3^{pS396}$ as in (A) normalised to respective total protein and WT DMSO-treated cells. Mean ± SD (n=3). Nd: not detected. Two-way ANOVA with Tukey's post-hoc comparison test.

**(C)** Confocal microscopy images of cells as in (A). Cyan: DNA (DAPI), grey: STING. Single Z-plane image, 40x oil immersion objective, scale bar = 10 µm.

**(D)** Relative mRNA levels of *IFNB1* in WT and *STING1* KO HMC3 cells treated with 10 nM AZD1390 (ATMi), or DMSO as a control, for 6 days. Cells were stimulated with 1 µM diABZI, or DMSO as a control, for 5 h. $C_q$ values normalised to *RPS13* and the DMSO-treated WT condition. Mean $\log_2$ fold change ± SD (n=3). Two-way ANOVA with Tukey's post-hoc comparison test.

**(E)** Concentration of interferons in cell culture supernatants. Mean ± SD (n=3). Dashed line indicates lower limit of detection (LLOD, pg/mL): IFN-α2a = 0.728, IFN-β = 3.28.

*p ≤ 0.05; **p ≤ 0.01; ****p ≤ 0.0001; ns = not significant.

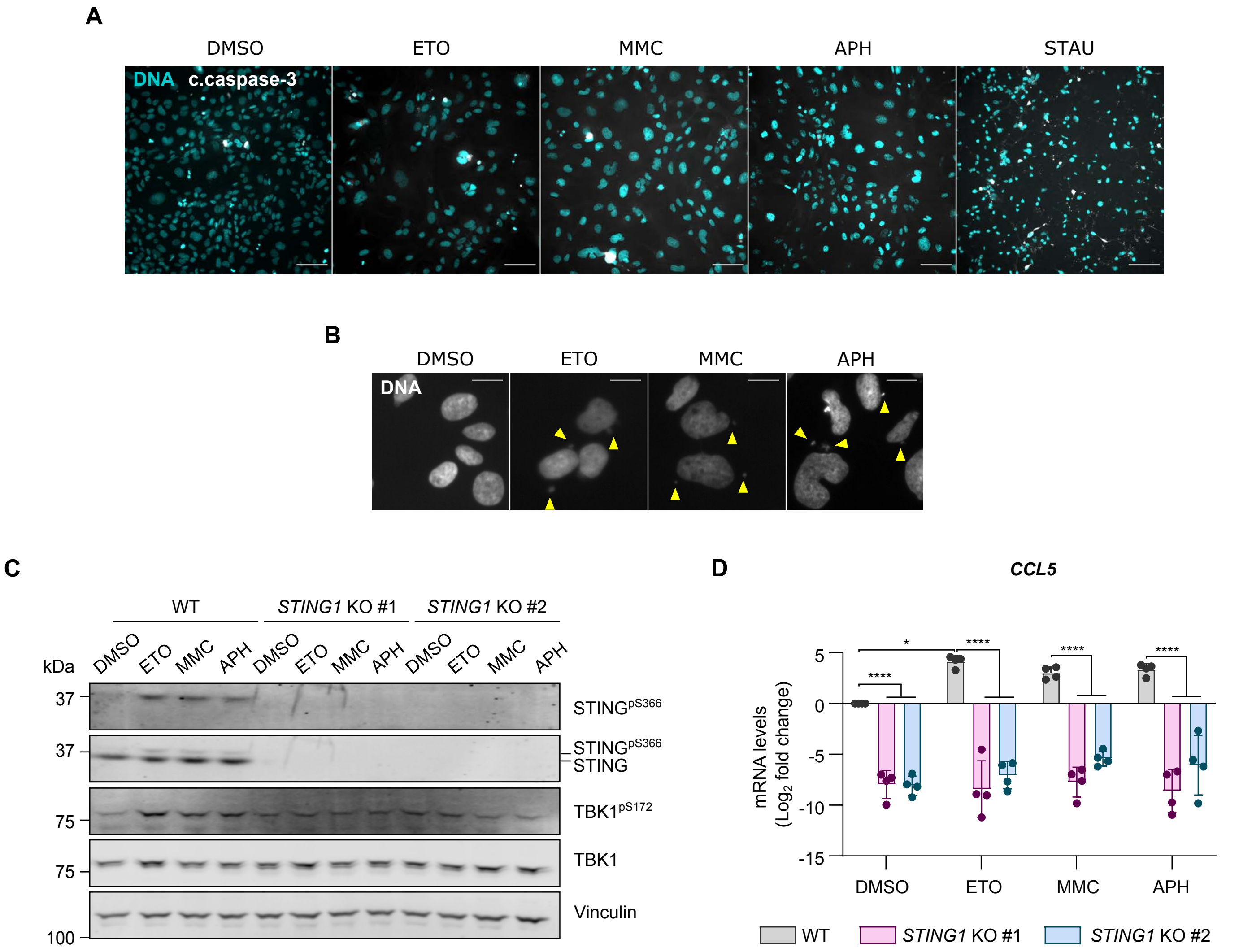


Talbot et al., Supplemental Figure 5

**Supplemental Figure 5. Persistent DNA damage drives STING-dependent chemokine induction in microglia-like cells.**

**(A)** Immunofluorescence microscopy images of WT and *STING1* KO HMC3 cells treated with 250 nM etoposide (ETO), 30 nM mitomycin C (MMC), 150 nM aphidicolin (APH), or DMSO as a control, for 6 days. Treatment with 1 µM staurosporine for 6 h served as a positive control of apoptosis. Cyan: DNA (DAPI), grey: cleaved caspase-3 (c.caspase-3). 20x objective, scale bar = 100 µm.
**(B)** Representative immunofluorescence microscopy images demonstrating the formation of micronuclei. Grey: DNA (DAPI). 20x objective, scale bar = 20 µm.
**(C)** Representative immunoblot of cells treated as in (A). Loading control: Vinculin.
**(D)** Relative mRNA levels of *CCL5*. $C_q$ values normalised to *RPS13* and the DMSO-treated WT condition. Mean $log_2$ fold change ± SD (n=3). Two-way ANOVA with Tukey's post-hoc comparison test.

*p ≤ 0.05; ****p ≤ 0.0001.

**Supplemental Table 1.** Cell lines, reagents and software programs used in this work.

| REAGENT or RESOURCE | SOURCE | IDENTIFIER |
|---|---|---|
| **Experimental Models: Cell Lines** | | |
| HMC3 cells | Kind gift from Dr Brian Bigger (University of Manchester, UK) | [1] |
| *STING1* knockout HMC3 cells | This paper | |
| C20 cells | Kind gift from Dr David Alvarez-Carbonell (Case Western Reserve University) | [2] |
| *ATM* knockout C20 cells | Khoronenkova Laboratory | [3] |
| iCell® Microglia | FujiFilm CDI | R1131 |
| **Bacterial Strains** | | |
| DH5α | Invitrogen | 18265017 |
| NEB® 10-beta | New England Biolabs | C3019 |
| **Recombinant DNA** | | |
| pSpCas9(BB)-2A-Puro (PX459) V2.0 | Kind gift from Feng Zhang | Addgene plasmid #62988; http://n2t.net/addgene:62988; RRID:Addgene_62988 [4] |
| **Oligonucleotides** | **Sequence (5' → 3')** | **SOURCE** |
| **Cloning** | | |
| STING sgRNA sense | caccgGCTGGGACTGCTGTTAAACG | This paper |
| STING1 sgRNA anti-sense | aaacCGTTTAACAGCAGTCCCAGCc | This paper |
| **Fragment analysis** | | |
| *STING1*-exon 3 Forward | GCTAGGCATCAAGGGAGTGA | This paper |
| *STING1*-exon 3 Reverse | TGGATTTCTTGGTGCCCACA | This paper |
| **RT-qPCR** | | |
| *CCL5* Forward | GCTGTCATCCTCATTGCTACTG | [5] |
| *CCL5* Reverse | TGGTGTAGAAATACTCCTTGATGTG | |
| *CGAS Forward* | CTGAACACCGGGAGCTACTA | This paper |
| *CGAS Reverse* | TTGAATTCTGGGGACTTCCAGT | |

| | | |
|---|---|---|
| *CXCL10* Forward | CCATTCTGATTTGCTGCCTTATC | [6] |
| *CXCL10* Reverse | TACTAATGCTGATGCAGGTACAG | |
| *IFIT2* Forward | CTGCAACCATGAGTGAGAACAA | This paper |
| *IFIT2* Reverse | CTCCCTCCATCAAGTTCCAGG | |
| *IFNB1* Forward | ATGACCAACAAGTGTCTCCTCC | [7] |
| *IFNB1* Reverse | GCTCATGGAAAGAGCTGTAGTG | |
| *IL6* Forward | AGACAGCCACTCACCTCTTCAG | [3] |
| *IL6* Reverse | TTCTGCCAGTGCCTCTTTGCTG | |
| *STING1* Forward | GGGCTGGCATGGTCATATTA | [8] |
| *STING1* Reverse | TACTCAGGTTATCAGGCACC | |
| *TFAM* Forward | GCGCTCCCCCTTCAGTTTTG | [9] |
| *TFAM* Reverse | GTTTTTGCATCTGGGTTCTGAGC | |
| *RPS13* Forward | CGAAAGCATCTTGAGAGGAACA | [3] |
| *RPS13* Reverse | TCGAGCCAAACGGTGAATC | |
| *IPO8* Forward | GGCACCACTCAGCGAGGAT | [10] |
| *IPO8* Reverse | TGTTGTTCAATCTTCTTCTTTGCCT | |
| **siRNA Oligonucleotides** | | |
| siCtrl (negative control) | GAUCGAUUCGCAGUUGAUUCU | This paper |
| sicGAS (SMART Pool) | Dharmacon siGENOME | M-015607-01 |
| siSTING | GGCAUCAAGGAUCGGGUUU | [11] |
| **Antibodies** (WB = western blotting; IF = immunofluorescence; FC = flow cytometry) | | |
| β-Actin (WB) | Abcam | ab6276 |
| ATM (WB) | Abcam | ab78 |
| ATM (WB) | Bethyl Laboratories | A300-299A |
| ATM pSer1981 (WB) | Abcam | ab81292 |
| BrdU (FC) | BD Biosciences | 347580 |
| cGAS (WB, IF) | Cell Signaling | 15102 |
| CHK2 (WB) | Cell Signaling | 3440 |
| CHK2 pThr68 (WB) | Cell Signaling | 2661 |
| Fibrillarin (WB) | Abcam | ab4566 |
| GAPDH (WB) | Cell Signaling | 2118 |
| GAPDH (WB) | Proteintech | 60004-1-Ig |

| | | |
|---|---|---|
| γH2AX (WB) | Millipore | 05-636 |
| IRF3 (WB) | Santa Cruz | sc-33641 |
| IRF3 (IF) | Cell Signaling | 11904 |
| IRF3 pSer396 (WB) | Cell Signaling | 4947 |
| STAT1 (WB) | Cell Signaling | 9176 |
| STAT1 pTyr701 (WB) | Cell Signaling | 9167 |
| STING (WB) | Cell Signaling | 13647 |
| STING (IF) | Abcam | ab181125 |
| STING pSer366 (WB) | Cell Signaling | 19781 |
| TBK1 (WB) | Abcam | ab40676 |
| TBK1 pSer172 (WB) | Cell Signaling | 5483 |
| α/β-Tubulin (WB) | Cell Signaling | 2148 |
| Vinculin (WB, IF) | Abcam | ab18058 |
| Vinculin (WB) | Proteintech | 26520-1-AP |
| AlexaFluor 680 goat anti-rabbit (WB) | Molecular Probes | A27042 |
| IR Dye 800 goat anti-mouse (WB) | LI-COR Biosciences | 926-32210 |
| AlexaFluor 488 goat anti-mouse (IF, FC) | Invitrogen | A11001 |
| AlexaFluor 488 goat anti-rabbit (IF) | Invitrogen | A32731 |
| AlexaFluor 594 goat anti-mouse (IF) | Invitrogen | A-11032 |
| Alexa Fluor 594 goat anti-rabbit (IF) | Abcam | ab150080 |
| **Chemicals, enzymes, and other reagents** | | |
| Aphidicolin | Merck | 178273 |
| AZD1390 (ATM inhibitor) | Selleckchem | S8680 |
| 5-Bromodeoxyuridine (BrdU) | Merck | B9285 |
| Camptothecin | Cambridge Bioscience | CAY11694 |
| RO-3306 (CDK1 inhibitor) | Tocris Bioscience | 4181 |
| Etoposide | Cambridge Bioscience | CAY12092 |
| Mitomycin C | Merck | M0503 |
| STING agonist diABZI | Selleckchem | S8796 |
| Puromycin | Santa Cruz | sc-108071 |

| | | |
|---|---|---|
| Recombinant human IFN-β | R&D Systems | 8499-IF |
| Phusion High-Fidelity DNA Polymerase | ThermoFisher | F530 |
| Lipofectamine RNAiMax | Invitrogen | 13778-150 |
| DNase I | ThermoFisher | EN0521 |
| 2-well silicone inserts | Ibidi | 80209 |
| TRI Reagent® | Merck | T9424 |
| **Kits and assays** | | |
| Glial-Mag Magnetofection | OzBiosciences | GL00500 |
| KAPA mRNA HyperPrep Kit | Roche | 08098115702 |
| NGS Fragment Kit (1-6,000 bp) | Agilent Technologies | DNF-473 |
| RNeasy Mini Kit | QIAGEN | 74106 |
| Agilent RNA 6000 Pico Kit | Agilent Technologies | 5067-1513 |
| RNAdvance Cell v2 | Beckman Coulter | A47943 |
| qPCRBIO cDNA Synthesis Kit | PCR Biosystems | PB30.11-10 |
| qPCRBIO SyGreen Blue Mix Lo-ROX | PCR Biosystems | PB20.15-20 |
| MSD U-PLEX human cytokine assay | Meso Scale Discovery | K151AEL-2 |
| **Software and Algorithms** | | |
| CellProfiler v4.0.7 | Broad Institute | |
| FlowJo v10.4.2 | FlowJo | |
| GraphPad Prism v9.4.1 | GraphPad Software | |
| Image Studio™ Lite Software v5.2 | LI-COR | |
| Image J | NIH | |
| DISCOVERY WORKBENCH v4.0 | Meso Scale Discovery | |
| QuantStudio™ Design & Analysis v1.5.1 | ThermoFisher | |

**Supplemental Table 2.** Human tissue sample information (source: NIH Neurobiobank, University of Maryland, United States). PMI = post-mortem interval, RIN = RNA integrity number, NA = not available.

| **ID** | **Diagnosis** | **Region** | **Brain Section** | **Age (years)** | **Sex** | **PMI (hours)** | **RIN** | **Cause of death** |
|---|---|---|---|---|---|---|---|---|
| 1259 | Control | Cerebellum | Vermis | 24 | Male | 8 | 9.2 | Accident MVA |
| 6018 | A-T | Cerebellum | Vermis | 24 | Male | 9 | 8.1 | Complication of Disorder |
| 5351 | Control | Cerebellum | Vermis | 44 | Male | 22 | 7 | Cardiac Arrhythmia |
| 5748 | A-T | Cerebellum | Vermis | 44 | Male | 29 | NA | Lymphoma / T Cell acute lymphoblastic leukemia |
| 6062 | Control | Cerebellum | Vermis | 32 | Female | 8 | 6 | Mixed Drug Intoxication |
| 6249 | A-T | Cerebellum | Vermis | 31 | Female | 4 | 3.2 | Complications of disorder |

### SUPPLEMENTAL REFERENCES


1. Janabi, N., Peudenier, S., Héron, B., Ng, K.H., and Tardieu, M. (1995). Establishment of human microglial cell lines after transfection of primary cultures of embryonic microglial cells with the SV40 large T antigen. Neurosci. Lett. *195*, 105–108. 10.1016/0304-3940(94)11792-H.

2. Garcia-Mesa, Y., Jay, T.R., Checkley, M.A., Luttge, B., Dobrowolski, C., Valadkhan, S., Landreth, G.E., Karn, J., and Alvarez-Carbonell, D. (2017). Immortalization of primary microglia: a new platform to study HIV regulation in the central nervous system. J. Neurovirol. *23*, 47–66. 10.1007/s13365-016-0499-3.

3. Bourseguin, J., Cheng, W., Talbot, E., Hardy, L., Lai, J., Jeffries, A.M., Lodato, M.A., Lee, E.A., and Khoronenkova, S.V. (2022). Persistent DNA damage associated with ATM kinase deficiency promotes microglial dysfunction. Nucleic Acids Res. *50*, 2700–2718. 10.1093/nar/gkac104.

4. Ran, F.A., Hsu, P.D., Wright, J., Agarwala, V., Scott, D.A., and Zhang, F. (2013). Genome engineering using the CRISPR-Cas9 system. Nat. Protoc. *8*, 2281–2308. 10.1038/nprot.2013.143.

5. Jenkins, R.W., Clarke, C.J., Canals, D., Snider, A.J., Gault, C.R., Heffernan-Stroud, L., Wu, B.X., Simbari, F., Roddy, P., Kitatani, K., et al. (2011). Regulation of CC Ligand 5/RANTES by Acid Sphingomyelinase and Acid Ceramidase. J. Biol. Chem. *286*, 13292–13303. 10.1074/jbc.M110.163378.

6. Lama, L., Adura, C., Xie, W., Tomita, D., Kamei, T., Kuryavyi, V., Gogakos, T., Steinberg, J.I., Miller, M., Ramos-Espiritu, L., et al. (2019). Development of human cGAS-specific small-molecule inhibitors for repression of dsDNA-triggered interferon expression. Nat. Commun. *10*, 2261. 10.1038/s41467-019-08620-4.

7. Kim, J.-E., Kim, Y.-E., Stinski, M.F., Ahn, J.-H., and Song, Y.-J. (2017). Human Cytomegalovirus IE2 86 kDa Protein Induces STING Degradation and Inhibits cGAMP-Mediated IFN-β Induction. Front. Microbiol. *8*, 154.

8. Wang, Y.-Y., Jin, R., Zhou, G.-P., and Xu, H.-G. (2016). Mechanisms of transcriptional activation of the stimulator of interferon genes by transcription factors CREB and c-Myc. Oncotarget *7*, 85049–85057. 10.18632/oncotarget.13183.

9. Zhang, R., and Wang, J. (2018). HuR stabilizes TFAM mRNA in an ATM/p38-dependent manner in ionizing irradiated cancer cells. Cancer Sci. *109*, 2446–2457. 10.1111/cas.13657.

10. Jansen, A.H., Mamber, C., Kooijman, L., Orre, M., Koot, S., Dooves, S., Dimayuga Smith, V., Kamphuis, W., Hol, E.M., Kirk, C.J., et al. (2013). Reactive glia show increased immunoproteasome activity in Alzheimer's disease. Brain *136*, 1415–1431. 10.1093/brain/awt083.

11. Herzner, A.-M., Hagmann, C.A., Goldeck, M., Wolter, S., Kübler, K., Wittmann, S., Gramberg, T., Andreeva, L., Hopfner, K.-P., Mertens, C., et al. (2015). Sequence-specific activation of the DNA sensor cGAS by Y-form DNA structures as found in primary HIV-1 cDNA. Nat. Immunol. *16*, 1025–1033. 10.1038/ni.3267.
